# Characterizing non-exponential growth and bimodal cell size distributions in Schizosaccharomyces pombe: an analytical approach

**DOI:** 10.1101/2021.06.10.447927

**Authors:** Chen Jia, Abhyudai Singh, Ramon Grima

## Abstract

Unlike many single-celled organisms, the growth of fission yeast cells within a cell cycle is not exponential. It is rather characterized by three distinct phases (elongation, septation and fission), each with a different growth rate. Experiments also show that the distribution of cell size in a lineage is often bimodal, unlike the unimodal distributions measured for the bacterium *Escherichia coli*. Here we construct a detailed stochastic model of cell size dynamics in fission yeast. The theory leads to analytic expressions for the cell size and the birth size distributions, and explains the origin of bimodality seen in experiments. In particular our theory shows that the left peak in the bimodal distribution is associated with cells in the elongation phase while the right peak is due to cells in the septation and fission phases. We show that the size control strategy, the variability in the added size during a cell cycle and the fraction of time spent in each of the three cell growth phases have a strong bearing on the shape of the cell size distribution. Furthermore we infer all the parameters of our model by matching the theoretical cell size and birth size distributions to those from experimental single cell time-course data for seven different growth conditions. Our method provides a much more accurate means of determining the cell size control strategy (timer, adder or sizer) than the standard method based on the slope of the best linear fit between the birth and division sizes. We also show that the variability in added size and the strength of cell size control of fission yeast depend weakly on the temperature but strongly on the culture medium.

**Author summary:** Advances in microscopy enable us to follow single cells over long timescales from which we can understand how their size varies with time and the nature of innate strategies developed to control cell size. This data shows that in many cell types growth is exponential and the distribution of cell sizes has one peak, namely there is a single characteristic cell size. However data for fission yeast shows remarkable differences: growth is non-exponential and the distribution of cell sizes has two peaks, meaning two characteristic cell sizes exist. Here we construct the first mathematical model of this organism; by solving the model analytically we show that it is able to predict the two peaked distributions of cell size seen in data and provides an explanation for each peak in terms of the various growth phases of the single-celled organism. Furthermore by fitting the model to the data, we infer values for the rates of all microscopic processes in our model. This method is shown to provide a much more reliable inference than current methods and sheds light on how the strategy used by fission yeast cells to control their size varies with external conditions.

## Introduction

Fission yeast cells are shaped as regular cylinders with hemispherical ends [1]. The cylinder has a fixed cross-sectional area and a variable length; hence both the length of the cylinder (cell length) and the area of the longitudinal section (cell area) are approximately proportional to cell volume. In experiments, length, area, and volume have all been used to characterize cell size. It has been reported that individual cell size grows exponentially in many cell types such as various bacterial strains and budding yeast [2–9]. However, fission yeast undergoes a complex non-exponential growth pattern within each cell cycle [10, 11], as illustrated by the time-course data of cell area along a typical cell lineage (Fig. 1(a)). Another remarkable feature of such lineage data is the bimodal shape of the cell size distribution (Fig. 1(b)). Recent studies have shown that if cell size grows exponentially in each generation, then the distribution of cell size must be unimodal [12, 13]. The main aim of the present paper is to propose a detailed model of cell size dynamics in fission yeast that can characterize its non-exponential growth, cell division, and size homeostasis, as well as develop an analytical theory that can account for the bimodal shape of the cell size distribution.

In the study of cell size dynamics, a core issue is to understand the size homeostasis strategies in various cell types, especially in fission yeast [14–17]. There are three popular phenomenological models of cell size control leading to size homeostasis [18]: (i) the timer strategy which implies a constant time between successive divisions; (ii) the sizer strategy which implies cell division upon attainment of a critical size, and (iii) the adder strategy which implies a constant size addition between consecutive generations. A conventional method of inferring the size control strategy is to use the information of cell sizes at birth and at division [19, 20]. This approach assumes that the birth size *V_b_* and the division size *V_d_* in each generation are related linearly by

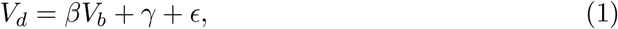

where 0 ≤ *β* ≤ 2 and *γ* ≥ 0 are two constants and *ϵ* is Gaussian white noise. Here *β* characterizes the strength of size control with *β* = 0, *β* =1, and *β* = 2 corresponding to the sizer, adder, and timer strategies, respectively. Using the data of birth and division sizes across different generations, the parameter *β* can be determined as the slope of the regression line of the division size on the birth size. However, in fission yeast, the linear relationship between birth and division sizes are actually very weak with numerous outliers and an exceptionally low *R*^2^ around 0.1 (Fig. 1(c)). This makes the inference of the parameter *β* highly unreliable. Hence another aim of the present paper is to develop a more reliable technique that can be used to accurately infer the size homeostasis strategy in fission yeast using a dynamic approach.

**Fig 1.**
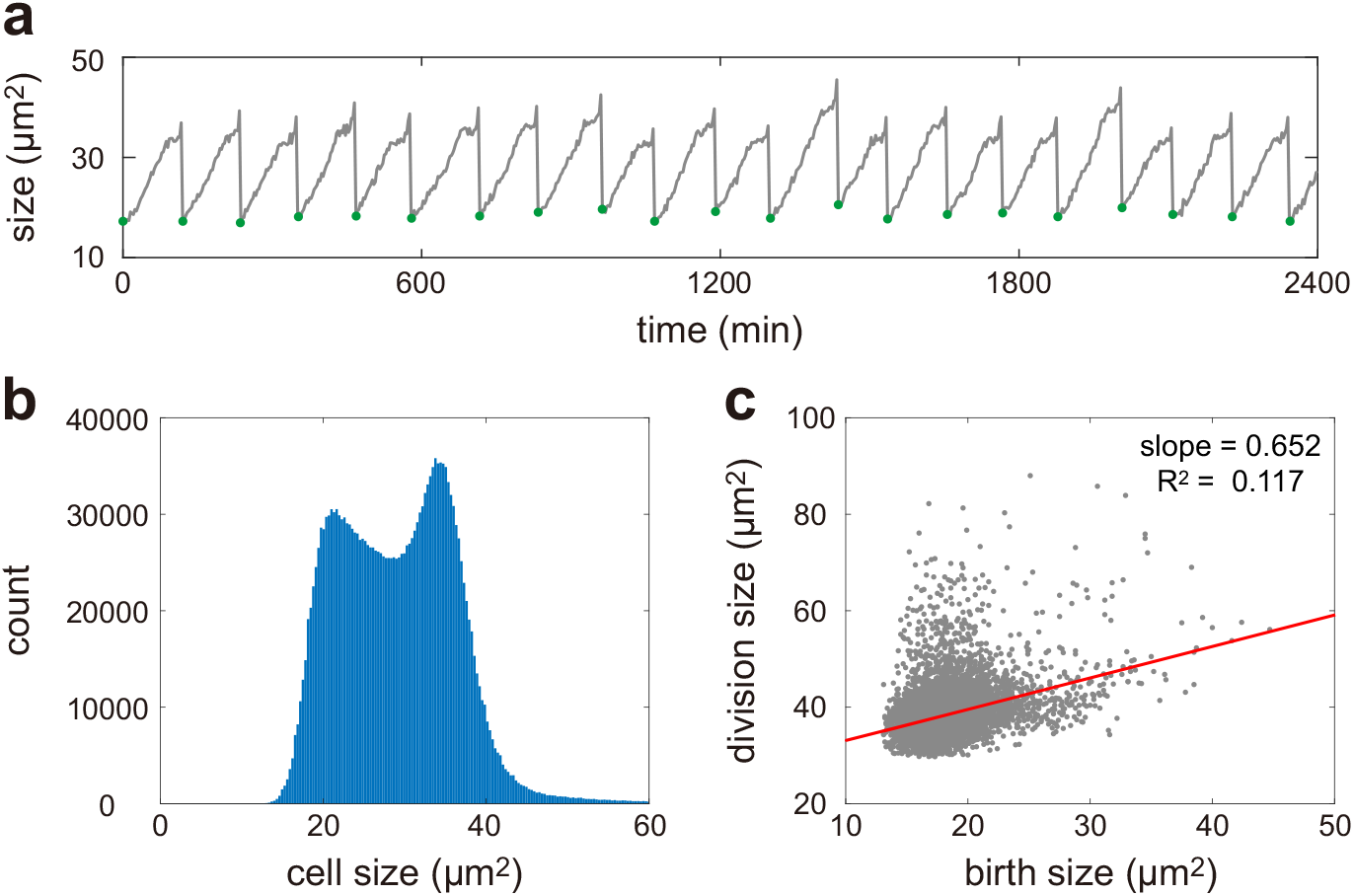
Cell size dynamics in fission yeast. (**a**) Single-cell time course data of cell area along a typical cell lineage cultured at 34°C in the yeast extract medium. The data shown are published in [11]. The green dots show cell sizes at birth. (**b**) Histogram of cell sizes along all cell lineages. The cell size distribution of lineage measurements has a bimodal shape. (**c**) Scatter plot of the birth size versus the division size and the regression line. When plotting (b),(c), we use the data of all 1500 cell lineages cultured at 34°C in the yeast extract medium, each of which is recorded every 3 minutes and is typically composed of 50 — 70 generations. The length of each generation is 114 ± 15 minutes.

## Results

### Model specification

Here we consider a detailed model of cell size dynamics in fission yeast across many generations, including a complex three-stage growth pattern, asymmetric and stochastic cell division, and size homeostasis (see Fig. 2(b) for an illustration). The model is based on a number of assumptions that are closely related to experimental data obtained using microfluidic devices. The assumptions are as follows and the specific meaning of all model parameters is listed in Table 1.

**Fig 2.**
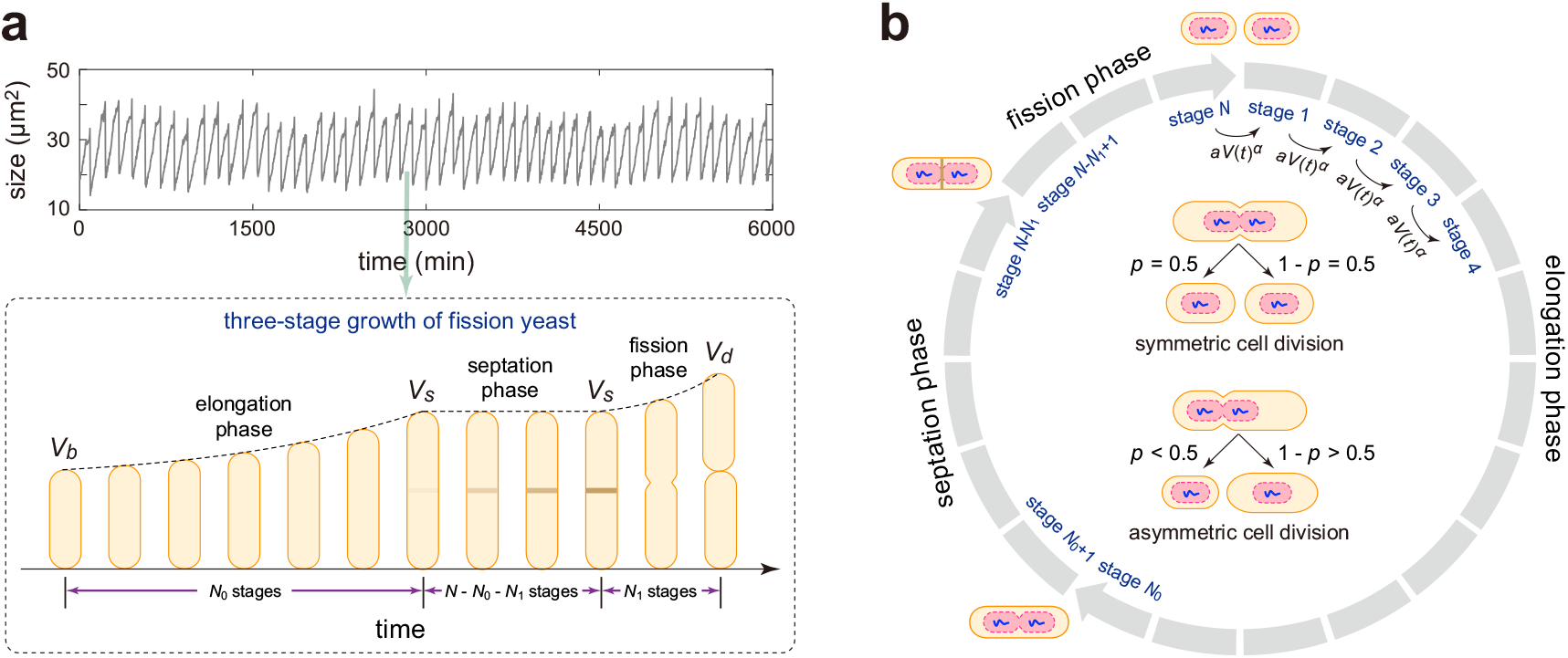
A detailed model of cell size dynamics in fission yeast. (**a**) Three-stage growth pattern of fission yeast: an elongation phase where cell size grows exponentially with rate *g*_0_, followed by a septation phase during which the septum is formed and cell size remains constant, and then followed by a fission phase where cell size increases abruptly with a higher growth rate *g*_1_ > *g*_0_. Here *V_b_* is the size at birth, *V_s_* is the size in the septation phase, and *V_d_* is the size at division. (**b**) Schematic illustrating a detailed model of cell size dynamics describing cell growth, multiple effective cell cycle stages, cell-size control, and symmetric or asymmetric partitioning at cell division (see inset graph). Each cell can exist in *N* effective cell cycle stages. The transition rate from one stage to the next at a particular time *t* is proportional to the *α*th power of the cell size *V*(*t*) with *α* > 0 being the strength of cell-size control and *a* > 0 being the proportionality constant. This guarantees that larger cells at birth divide faster than smaller ones to achieve size homeostasis. At stage *N*, a mother cell divides into two daughters that are typically different in size via asymmetric cell division. Symmetric division is the special case where daughters are equisized.

#### 1)

The growth of cell size of fission yeast within a cell cycle is very different from the exponential growth observed in many other cell types [2]. Actually, fission yeast undergoes a non-exponential three-stage growth pattern: an elongation phase followed by a septation phase and a fission phase (Fig. 2(a)) [10]. During the elongation phase, the size of each cell grows exponentially in each generation with growth rate *g*_0_. Note that in some previous papers [10], the growth in the elongation phase is assumed to be linear. However, numerous single-cell time-course measurements of cell size under different growth conditions support the assumption of exponential elongation used in the present paper [11]. After the elongation phase, the size of the cell remains constant for a period during which the septum is formed (see Fig. 4A in [10]). From the lineage data in Fig. 1(a), it seems also reasonable to assume that there is a small non-zero growth rate in the septation phase. However, according to the principle of parsimony, we choose not to introduce an extra parameter and assume zero growth rate during septation. At the end of the septation phase, there is a sharp increase in cell size for a short period prior to division; this period is called the fission phase. During the fission phase, we assume that cell size grows exponentially with a higher rate *g*_1_ > *g*_0_.

**Table 1.** Model parameters and their meaning.

| model parameters | meaning |
| --- | --- |
| $g_0$ | exponential growth rate in the elongation phase |
| $g_1$ | exponential growth rate in the fission phase |
| $N$ | total number of effective cell cycle stages |
| $N_0$ | number of cell cycle stages in the elongation phase |
| $N_1$ | number of cell cycle stages in the fission phase |
| $r_0$ | proportion of cell cycle stages in the elongation phase |
| $r_1$ | proportion of cell cycle stages in the fission phase |
| $a$ | proportionality constant for the transition rate between stages |
| $\alpha$ | strength of size control |
| $M_0$ | mean generalized added size in the elongation phase |
| $M_1$ | mean generalized added size in the fission phase |
| $p$ | mean partition ratio of cell size at division |

#### 2)

Each cell can exist in *N* effective cell cycle stages, denoted by 1, 2,…, *N*. We assume that the cell stays in the elongation phase in the first *N*_0_ stages, stays in the fission phase in the last *N*_1_ stages, and stays in the septation phase in the intermediate *N* – *N*_0_ – *N*_1_ stages (Fig. 2). The transition rate from one stage to the next at a particular time is proportional to the *α*th power of cell size at that time [13, 21]. In other words, the transition rate between stages at time t is equal to *aV*(*t*)^*α*^, where *V*(*t*) is the cell size at that time, *α* > 0 is the strength of cell size control, and *a* > 0 is a proportionality constant. Under this assumption, larger cells at birth, on average, have shorter cell cycle duration and lesser volume change than smaller ones; in the way size homeostasis is achieved.

Recent investigations have suggested that the accumulation of some cyclin (Cdc13), phosphatase (Cdc25), or kinase (Cdr2) up to a critical threshold as a possible mechanism for fission yeast cell division [14–16]. Biophysically, the *N* effective cell cycle stages in our model can be understood as different levels of the division protein (Cdc13, Cdc25, or Cdr2). The power law form for the rate of cell cycle progression may come from cooperation of the division protein, as explained in detail in [13, 21]. This power law not only coincides with certain biophysical mechanisms, but also results in a natural scaling transformation among the timer, sizer, and adder, as will be explained later.

Let *V_b_* and *V_d_* denote cell sizes at birth and at division in a particular generation, respectively, and let *V_s_* denote cell size in the septation phase, which is assumed to be a constant. Then the increment in the *α*th power of cell size, which is referred to as generalized added size, in the elongation phase, 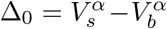, has an Erlang distribution with shape parameter *N*_0_ and mean *M*_0_ = *N*_0_*g*_0_*α/a* (see Supplementary Section 1 for the proof). Similarly, the generalized added size in the fission phase, 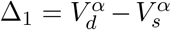, also has an Erlang distribution with shape parameter *N*_1_ and mean *M*_1_ = *N*_1_*g*_1_*α/a*. Therefore, the total generalized added size across the cell cycle, 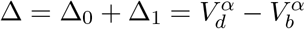, is the sum of two independent Erlang distributed random variables and has a hypoexponential distribution (also called generalized Erlang distribution) whose Laplace transform is given by

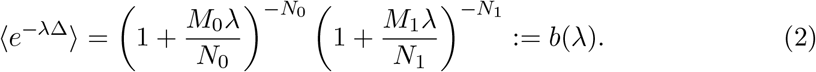

Note that 〈*e*^−λΔ^〉 → *e*^(*M*_0_+*M*_1_)λ^ as *N* → ∞. This means that the generalized added size Δ = *M*_0_ + *M*_1_ becomes deterministic when *N* is large. However, when *N* is small, the variability in Δ is much larger. Hence, our model allows the investigation of the influence of added size variability on cell size dynamics.

Three special cases deserve special attention. When *α* → 0, the transition rate between stages is a constant and thus the doubling time has an Erlang distribution that is independent of the birth size; this corresponds to the timer strategy. When *α* = 1, the added size *V_d_* – *V_b_* has an hypoexponential distribution that is independent of the birth size; this corresponds to the adder strategy. When *α* → ∞, the *α*th power of the division size, 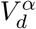, has a hypoexponential distribution that is independent of the birth size; this corresponds to the sizer strategy. Intermediate strategies are naturally obtained for intermediate values of *α*; timer-like control is obtained when 0 < *α* < 1 and sizer-like control is obtained when 1 < *α* < ∞ [21].

#### 3)

Cell division occurs when the cell transitions from stage *N* to the next stage 1. At division, most previous papers assume that the mother cell divides into two daughters that are exactly the same in size via symmetric partitioning [22–25]. Experimentally, fission yeast in general do not divide perfectly in half. Here we follow the methodology that we devised in [13, 26] and extend previous models by considering asymmetric partitioning at division: the mother cell divides into two daughters with different sizes.

If the partitioning of cell size is symmetric, we track one of the two daughters randomly after division [27, 28]; if the partitioning is asymmetric, we either track the smaller or the larger daughter after division [29, 30]. Let *V_d_* and 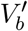 denote cell sizes at division and just after division, respectively. If the partitioning is deterministic, then we have 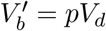, where 0 < *p* < 1 is a constant with *p* = 0.5 corresponding to symmetric division, p < 0.5 corresponding to smaller daughter tracking, and *p* > 0.5 corresponding to larger daughter tracking. The value of *p* can be inferred from experiments. However, in fission yeast, the partitioning of cell size is appreciably stochastic. In this case, we assume that the partition ratio 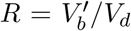 has a beta distribution with mean *p* [31], whose probability density function is given by

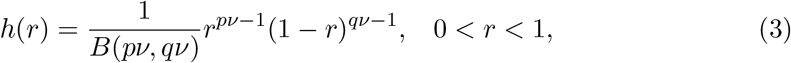

where *B* is the beta function, *q* = 1 – *p*, and *ν* > 0 is referred to as the sample size parameter. When *ν* → ∞, the variance of the beta distribution tends to zero and thus stochastic partitioning reduces to deterministic partitioning, i.e. *f*(*r*) = *δ*(*r* – *p*).

We next describe our stochastic model of cell size dynamics. The microstate of the cell can be represented by an ordered pair (*k,x*), where *k* is the effective cell cycle stage which is a discrete variable and *x* is the cell size which is a continuous variable. Note that the cell undergoes deterministic growth in each stage (exponential growth in the first *N*_0_ and the last *N*_1_ stages and no growth in the remaining *N* – *N*_0_ – *N*_1_ stages), and the system can hop between successive stages stochastically. Let *p_k_*(*x*) denote the probability density function of cell size when the cell is in stage *k*. Then the evolution of cell size dynamics in fission yeast can be described by a piecewise deterministic Markov process whose master equation is given by

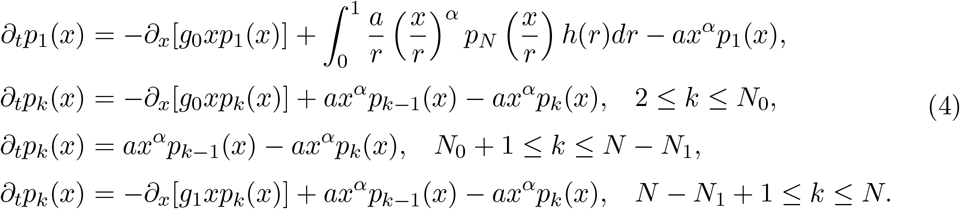

where *h*(*r*) is the function given in Eq. (3). In the first, second, and fourth equations, the first term on the right-hand side describe cell growth and the remaining two terms describe transitions between cell cycle stages. In the third equation, the two terms on the right-hand side describe cell cycle stage transitions. In the first equation, the middle term on the right-hand side describes the partitioning of cell size at division.

### Analytical distribution of cell size of lineage measurements

Let 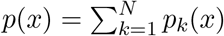 denote the probability density function of cell size *V* (here we use *V* to represent a random variable and use *x* to represent a realization of *V*). In our model, we assume that the rate of cell cycle progression has a power law dependence on cell size. This assumption implies an important scaling property of our model: if the dynamics for cell size *V* has a control strength *α* (with *α* < 1 corresponding to timer-like and *α* > 1 corresponding to sizer-like strategies), then the dynamics for the *α*th power of cell size, *V^α^*, has an adder strategy. This scaling property serves as the key to our analytical theory.

Recall that the probability distribution of any random variable with nonnegative values is fully determined by its Laplace transform. To obtain the analytical distribution of cell size along a cell lineage, we introduce 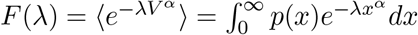, which is nothing but the Laplace transform for the *α*th power of cell size. For simplicity, we first focus on the case of deterministic partitioning. Despite the biological complexity described by our model, the Laplace transform can still be solved exactly in steady-state conditions as (see Supplementary Section 2 for the proof)

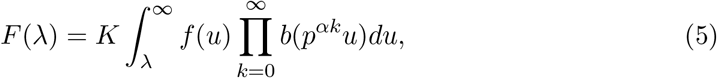

where *b*(λ) is the function given in Eq. (2),

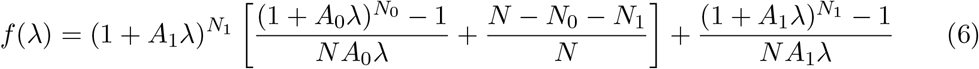

is another function with *A*_0_ = *M*_0_/*N*_0_ and *A*_1_ = *M*_1_/*N*_1_, and

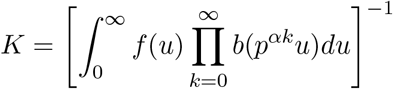

is a normalization constant. From the definition of *f*(λ) in Eq. (6), it is clear that *f*(λ) tends to infinity as λ → ∞. However, from the definition of *b*(λ) in Eq. (2), the infinite product 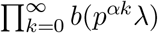 decays to zero as λ → ∞ at a faster exponential speed. Hence the integral in Eq. (5) is always well defined.

In principle, taking the inverse Laplace transform gives the probability density function of *V^α^*, from which the distribution of cell size *V* can be obtained. Next we introduce how to compute the cell size distribution more effectively using our analytical results. Taking the derivative with respect to λ on both sides of Eq. (5), using the change of variables formula, and finally replacing λ by *i*λ yield (see Supplementary Section 2 for the proof)

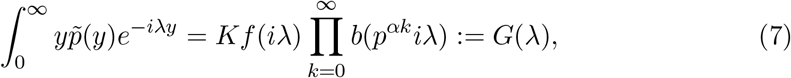

where

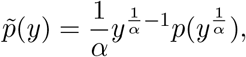

is the probability density function of *V^α^*. This shows that the Fourier transform of 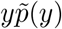 is exactly *G*(λ). Since the Fourier transform and inverse Fourier transform are inverses of each other, we only need to take the inverse Fourier transform of *G*(λ) so that we can obtain 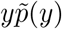. Finally, the cell size distribution *p*(*x*) can be recovered from 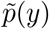 as

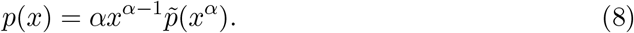

In general, the cell size distribution along a cell lineage can also be numerically computed by carrying out stochastic simulations of the piecewise deterministic Markovian model. However, under the complex three-stage growth pattern of fission yeast, according to our simulations, over 10^7^ stochastic trajectories must be generated in order to obtain an accurate computation of the size distribution (Supplementary Fig. 1) — this turns out to be very slow. The analytical solution is thus important since it allows a fast exploration of large swathes of parameter space without performing stochastic simulations.

To gain deeper insights into the cell size distribution, we next consider two important special cases. For the case of exponential growth of cell size, there is only the elongation phase and the remaining two phases vanish. In this case, we have *N*_1_ = 0 and *N* = *N*_0_; the cell size distribution is still determined by Eq. (5) with the functions *b*(λ) and *f*(λ) being simplified greatly as

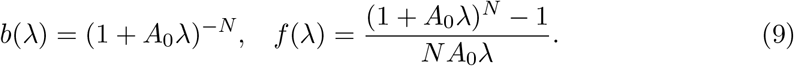

In fact, the analytical cell size distribution for exponentially growing cell lineages has been studied previously in [13], where the distribution of the logarithmic cell size, instead of the original cell size, is obtained. We emphasize that the analytical expression given here is not only much simpler, but also numerically more accurate than the one given in that paper, which includes the integral of an infinite product term which is very difficult to compute accurately.

The second case occurs when *N* → ∞, while keeping *r*_0_ = *N*_0_/*N* and *r*_1_ = *N*_1_/*n* as constants, where *r*_0_ and *r*_1_ represent the proportions of cell cycle stages in the elongation and fission phases, respectively. In this case, the generalized added size Δ becomes deterministic and the system does not involve any stochasticity. As *N* → ∞, the Laplace transform given in Eq. (5) can be simplified to a large extent as (see Supplementary Section 2 for the proof)

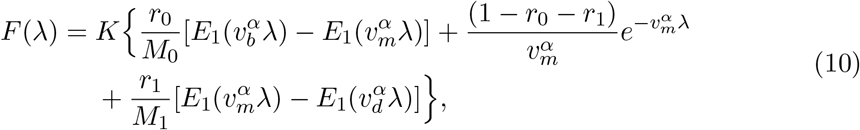

where 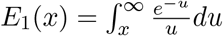 is the exponential integral,

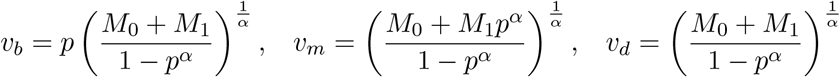

are the birth size, septation size, and division size, respectively, and *K* = (*T*_1_ + *T*_2_ + *T*_3_)^−1^ is a normalization constant with

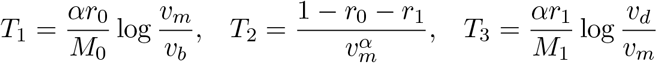

being the durations in the elongation, septation, and fission phases, respectively. Note that in [10], the septation size (size in the septation phase) is called the division size and the division size (size just before division) is called the fission size. Here the terminology is slightly different. Taking the inverse Laplace transform finally gives the cell size distribution:

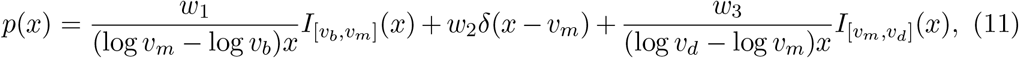

where *I_A_*(*x*) is the indicator function which takes the value of 1 when *x* ∈ *A* and takes the value of 0 otherwise, *δ*(*x*) is Dirac’s delta function, and

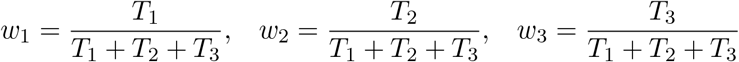

are the proportions of subpopulations in the three phases, respectively. This indicates that when added size variability is small, cell size has a distribution that is concentrated on a finite interval between *v_b_* and *v_d_*.

To validate our theory, we compare the analytical cell size distribution with the one obtained from stochastic simulations under different choices of *N* (Fig. 3(a)). Clearly, they coincide perfectly with each other. It can be seen that as added size variability become smaller (*N* increases), the analytical distribution given in Eq. (8) converges to the limit distribution given in Eq. (11). When *N* is small, the size distribution is unimodal. As *N* increases, the size distribution becomes bimodal with the right peak becoming higher and narrower. The bimodality of the size distribution can be attributed to cells in different phases: *the left peak corresponds to cells in the elongation phase and the right peak corresponds to cells in the septation and fission phases.* Since the size in the elongation phase is smaller than that in the fission or septation phase, the left peak is associated with elongation and the right peak with the other two phases. When *N* is very large, the size distribution is the superposition of three terms, corresponding to the three phases of cell growth.

To gain a deeper insight, we illustrate the cell size distribution as a function of the parameters *α*, *r*_0_, *r*_1_, and *g*_1_ when *N* is relatively large (Fig. 3(b)-(e)). It can be seen that as size control becomes stronger (*α* increases), the size distribution changes from the unimodal to the bimodal shape (Fig. 3(b)). The size distribution is generally unimodal for timer-like strategies and bimodal for sizer-like strategies. The dependence of the size distribution on *r*_0_ is expected — a small *r*_0_ results in a small fraction of cells in the elongation phase and thus the left peak is much lower than the right peak, while a large *r*_0_ gives rise to the opposite effect (Fig. 3(c)). Bimodality is the most apparent when *r*_0_ is neither too large nor too small.

**Fig 3.**
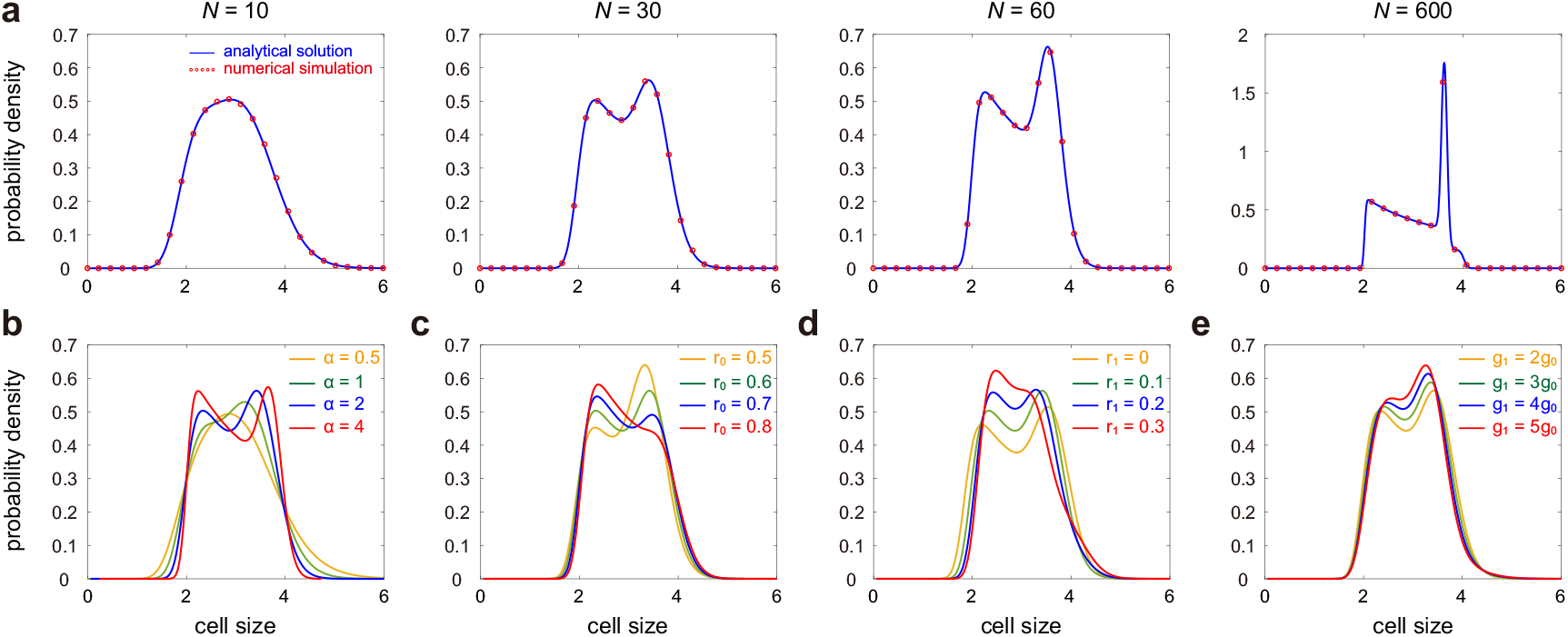
Influence of model parameters on the cell size distribution. (**a**) Cell size distribution as *N* varies. The blue curve shows the analytical distribution obtained by taking the inverse Laplace transform of Eq. (5) (e.g. by using the technique described by Eqs. (7) and (8)) and the red circles show the distribution obtained using stochastic simulations. The parameters are chosen as *r*_0_ = 0.6, *r*_1_ = 0.1, *g*_1_ = 2*g*_0_, *α* = 2. (**b**) Cell size distribution as *α* varies. The parameters are chosen as *N* = 30, *r*_0_ = 0.6, *r*_1_ = 0.1, *g*_1_ = 2*g*_0_. (**c**) Cell size distribution as *r*_0_ varies. The parameters are chosen as *N* = 30, *r*_1_ = 0.1, *g*_1_ = 2*g*_0_, *α* = 2. (**d**) Cell size distribution as *r*_1_ varies. The parameters are chosen as *N* = 30, *r*_0_ = 0.6, *g*_1_ = 2*g*_0_, *α* = 2. (**e**) Cell size distribution as *g*_1_ / *g*_0_ varies. The parameters are chosen as *N* = 30, *r*_0_ = 0.6, *r*_1_ = 0.1, *α* = 2. In (a)-(e), the parameters *g*_0_ and *p* are chosen as *g*_0_ = 0.01, *p* = 0.5 and the parameters a, *M*_0_, *M*_1_ are chosen so that the mean cell size 〈*V*〉 = 3.

The influence of *r*_1_ on the cell size distribution is more complicated. Recall that a larger *r*_1_ means a larger fraction of cells in the fission phase and a smaller fraction of cells in the septation phase. Here since we fix *r*_0_ to be a constant and tune *r*_1_, there is little change in the fraction of cells in the elongation phase. As the septation phase becomes shorter (*r*_1_ increases), the size distribution changes from being bimodal to being unimodal and becomes more concentrated (Fig. 3(d)). In particular, bimodality is apparent when the septation phase is relatively long, while a very short septation phase may even destroy bimodality.

Finally, we examine the dependence of the cell size distribution on the ratio of the growth rate in the fission phase to the one in the elongation phase, *g*_1_/*g*_0_, which characterizes the sharpness of the size increase in the fission phase. As the size addition in the fission phase becomes sharper (*g*_1_/*g*_0_ increases), the size distribution changes from being bimodal to being unimodal and becomes more concentrated (Fig. 3(d)). Here we fix the mean cell size to be a constant by tuning the parameter a and thus the increase in *g*_1_ does not make the right peak shift more to the right. To our surprise, we find that bimodality is the most apparent when the growth rates in the two phases are close to each other, while a very abrupt size addition in the fission phase may even destroy bimodality.

To summarize, we find that small added size variability, strong size control, moderate length in the elongation phase, long septation phase, short fission phase, and mild size addition in the fission phase are capable of producing more apparent bimodality.

### Analytical distribution of the birth size

In our model, the distribution of the birth size *V_b_* can also be derived analytically in steady-state conditions. In fact, the Laplace transform for the *α*th power of the birth size, 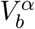, is given by (see Supplementary Section 3 for the proof)

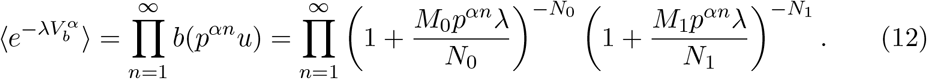

Taking the inverse Laplace transform gives the probability density function of 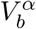, from which is the distribution of *V_b_* can be obtained. A special case takes place when *α* is large (strong cell-size control) or when *p* is small (smaller daughter tracking). Under the large *α* or small *p* approximation, the term *p^αn^* is negligible for *n* ≥ 2 and it suffices to keep only the first term in the infinite product given in Eq. (12). In this case, the laplace transform of 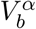 reduces to

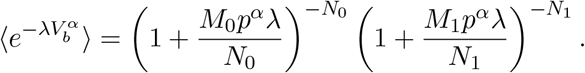

Taking the inverse Laplace transform gives the birth size distribution

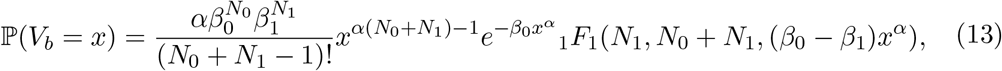

where _1_*F*_1_ is the confluent hypergeometric function, *β*_0_ = *N*_0_/*M*_o_*p*^*α*^, and *β*_0_ = *N*_1_/*M*_1_*p*^*α*^.

Actually, the birth size distribution has also been computed analytically in some simpler models. It has been shown that the birth size in those models approximately has a log-normal distribution [22] or a gamma distribution [32]. Therefore it is natural to ask whether the birth size in our model shares the same property. To see this, we illustrate the birth size distribution and its approximation by the log-normal and gamma distributions as *N* and *α* vary (Fig. 4(a),(b))). We find that under a wide range of model parameters, the true distribution is in excellent agreement with its log-normal approximation. However, when *N* and *α* are both small, the true distribution is severely right-skewed and deviates significantly from its gamma approximation. When *N* and *α* are both large, the true distribution becomes more symmetric and the three distributions become almost indistinguishable.

**Fig 4.**
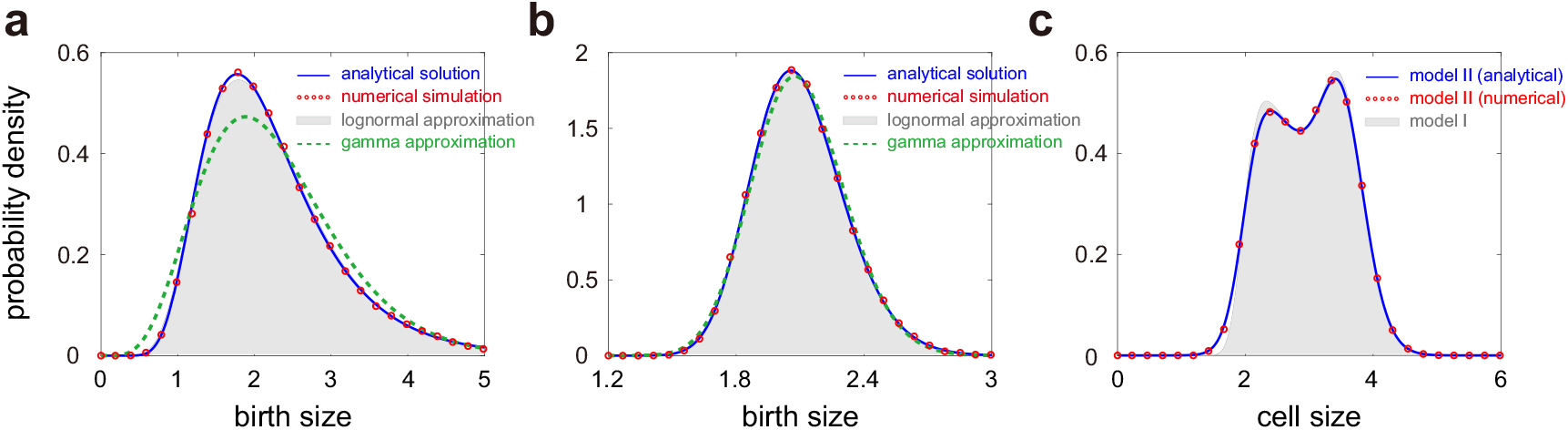
Further properties of the birth size and cell size distributions. (**a**) Comparison of the birth size distribution (blue curve and red circles) with its log-normal (solid grey region) and gamma (dashed green curve) approximations when *N* and *α* are small. The blue curve shows the analytical distribution obtained by taking the inverse Laplace transform of Eq. (12) and the red circles show the distribution obtained from stochastic simulations. (**b**) Same as (a) but when *N* and *α* are large. In (a),(b), the parameters are chosen as *r*_0_ = 0.6, *r*_1_ = 0.1, *g*_0_ = 0.01, *g*_1_ = 4*g*_0_, *p* = 0.5. The parameters *N* and *α* are chosen as *N* = 10, *α* = 0.5 in (a) and *N* = 30, *α* = 2 in (b). (**c**) Comparison between the cell size distributions for the model with deterministic partitioning (solid grey region) and the model with stochastic partitioning (blue curve and red circles). The blue curve shows the analytical distribution obtained by taking the inverse Laplace transform of Eq. (14) and the red circles show the distribution obtained from stochastic simulations. The parameters are chosen as *N* = 30, *r*_0_ = 0.6, *r*_1_ = 0.1, *g*_0_ = 0.01, *g*_1_ = 4*g*_0_, *α* = 2, *p* = 0.5. For the model with stochastic partitioning, the parameter *ν* is chosen as *v* = 200. In (a)-(c), the parameters a, *M*_0_, *M*_1_ are chosen so that the mean cell size 〈*V*〉 = 3 for the model with deterministic partitioning.

### Influence of stochastic partitioning on the cell size distribution

Thus far, the analytical distribution of cell size is derived when the partitioning at division is deterministic. In the presence of noise in partitioning, we can also obtain an explicit expression for the cell size distribution, whose Laplace transform is given by (see Supplementary Section 2 for the proof)

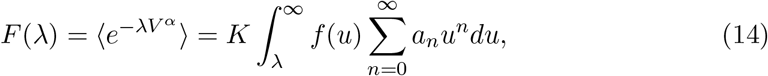

where *f*(λ) is the function given in Eq. (6),

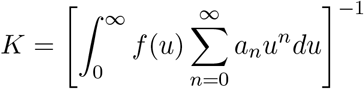

is a normalization constant, and *a_n_* is a sequence that can be determined by the following recursive relations:

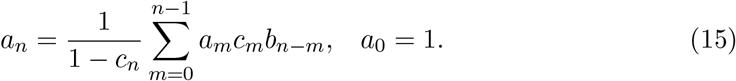

Here *b_n_* and *c_n_* are two other sequences that are defined by

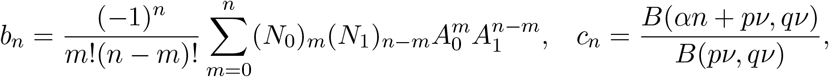

with (*x*)_*m*_ = *x*(*x* + 1) ⋯ (*x* + *m* – 1) being the Pochhammer symbol. For the special case of exponential growth of cell size, there is only the elongation phase and the remaining two phases vanish. In this case, we have *N*_1_ = 0 and *N* = *N*_0_; the cell size distribution is still determined by Eq. (14) with the sequence *b_n_* and the function *f*(λ) being greatly simplified as

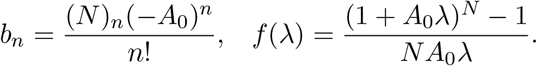

Clearly, fluctuations in partitioning at division lead to a much more complicated analytical expression of the cell size distribution. Actually, when partitioning is stochastic, the analytical cell size distribution for exponentially growing cell lineages has been obtained approximately in [13] under the assumption that noise in partitioning is very small. Here we have removed this assumption and obtained a closed-form solution of the size distribution for general non-exponentially growing cell lineages even if noise in partitioning is very large. Recent cell lineage measurements suggest that the coefficient of variation of the partition ratio 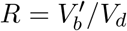 in fission yeast is 6% - 8% under different growth conditions [11].

To see the effect of stochastic partitioning, we illustrate the cell size distributions under deterministic and stochastic partitioning in Fig. 4(c) with the standard deviation of the partition ratio *R* being 7% of the mean for the latter. Clearly, the analytical solution given in Eq. (14) matches the simulation results very well. In addition, it can be seen that noise in partitioning gives rise to larger fluctuations in cell size, characterized by a smaller slope of the left shoulder, an apparent decrease in the height of the left peak, and a slight decrease in the height of the right peak. The valley between the two peaks and the right shoulder are almost the same for the two models.

### Correlation between sizes at birth and sizes at division

In [22], it has been shown that the correlation between cell sizes at birth and at division can be used to infer the size control strategy. For the case of deterministic partitioning, since the generalized added size 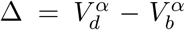 is hypoexponentially distributed, it is easy to obtain (see Supplementary Section 4 for the proof)

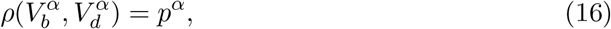

where *ρ*(*X, Y*) denotes the correlation coefficient between *X* and *Y*. This characterizes the correlation between sizes at birth and sizes at division in fission yeast, which only depends on the asymmetry of partitioning (*p*) and the strength of size control (*α*). In particular, we find that if partitioning is deterministic, the correlation between birth and division sizes is independent of the growth pattern of the cell — both exponentially and non-exponentially growing cells share the same correlation coefficient whenever they are the same parameters *p* and *α*.

In the presence of noise in partitioning, the formula for the correlation coefficient should be modified as (see Supplementary Section 4 for the proof)

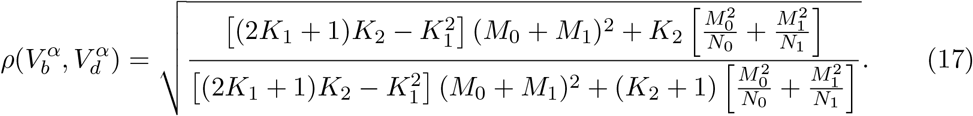

where

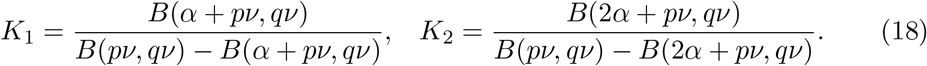

In this case, 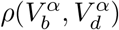 is generally lower than *p^α^* due to partitioning noise. Interestingly, if partitioning is stochastic, the correlation between birth and division sizes not only depends on *p* and *α*, but also depends on the parameters *N*_0_, *M*_0_, *N*_1_, *M*_1_, which describe the growth pattern of fission yeast. This is very different from the case of deterministic partitioning.

### Experimental validation of the theory

To test our theory, we apply it to lineage data of cell size in fission yeast that are published in [11]. In this data set, the single-cell time-course data of cell size were recorded every three minutes using microfluidic devices. The experiments were performed under seven growth conditions with different media (Edinburgh minimal medium (EMM) and yeast extract medium (YE)) and different temperatures. For EMM, cells were cultured at four different temperatures (28°C, 30°C, 32°C, and 34°C), while for YE, three different temperatures (28°C, 30°C, and 34°C) were used. For each growth condition, 1500 cell lineages were tracked and each lineage is typically composed of 50-70 generations. Note that for a particular cell lineage, it may occur that the cell was dead or disappeared from the channel during the measurement [11, 33]. Such lineages are removed from the data set and thus the number of lineages used for data analysis for each growth condition is actually less than 1500.

Based on such data, it is possible to estimate all the parameters involved in our model for the seven growth conditions. Parameter inference is crucial since it provides insights into the size control strategy, added size variability, and complex growth pattern in fission yeast. We perform parameter inference by fitting the noisy data to two models: the model with deterministic partitioning (model I) and the model with stochastic partitioning (model II). The estimated values of all parameters for the two models are listed in Table 2. Compared with model II, model I gives rise to a lower estimate of *α* and *r*_1_, as well as a higher estimate of *N*, *a*, and *g*_1_; the estimates of other parameters are very similar for the two models. In the following, we briefly describe our parameter estimation method.

**Table 2.** Parameters estimated using lineage data of cell size in fission yeast under seven growth conditions. Two theoretical models are used: the model with deterministic partitioning (model I) and the model with stochastic partitioning (model II). The estimation error for each parameter was computed using bootstrap. Specifically, we performed parameter inference 50 times; for each estimation, the theoretical model was fitted to the data of 50 randomly selected cell lineages. The estimation error was then calculated as the standard deviation over 50 repeated samplings.

| model I | EMM 28°C | EMM 30°C | EMM 32°C | EMM 34°C | YE 28°C | YE 30°C | YE 34°C |
| --- | --- | --- | --- | --- | --- | --- | --- |
| $p$ | $0.459 \pm 0.003$ | $0.459 \pm 0.002$ | $0.468 \pm 0.003$ | $0.470 \pm 0.004$ | $0.466 \pm 0.003$ | $0.468 \pm 0.003$ | $0.475 \pm 0.004$ |
| $\alpha$ | $1.767 \pm 0.093$ | $1.695 \pm 0.089$ | $1.726 \pm 0.102$ | $1.692 \pm 0.097$ | $1.139 \pm 0.058$ | $1.371 \pm 0.072$ | $1.245 \pm 0.066$ |
| $N$ | $17.463 \pm 0.803$ | $20.727 \pm 0.899$ | $20.002 \pm 0.886$ | $21.010 \pm 0.900$ | $32.369 \pm 1.375$ | $45.713 \pm 1.846$ | $55.315 \pm 2.126$ |
| $r_0$ | $0.645 \pm 0.011$ | $0.658 \pm 0.015$ | $0.654 \pm 0.012$ | $0.696 \pm 0.018$ | $0.709 \pm 0.018$ | $0.651 \pm 0.012$ | $0.634 \pm 0.010$ |
| $r_1$ | $0.070 \pm 0.006$ | $0.039 \pm 0.004$ | $0.045 \pm 0.004$ | $0.034 \pm 0.003$ | $0.038 \pm 0.004$ | $0.041 \pm 0.005$ | $0.034 \pm 0.004$ |
| $a$ | $0.012 \pm 0.001$ | $0.022 \pm 0.003$ | $0.019 \pm 0.002$ | $0.022 \pm 0.003$ | $0.301 \pm 0.035$ | $0.210 \pm 0.029$ | $0.420 \pm 0.051$ |
| $g_0$ | $0.216 \pm 0.001$ | $0.277 \pm 0.002$ | $0.279 \pm 0.002$ | $0.245 \pm 0.002$ | $0.333 \pm 0.003$ | $0.400 \pm 0.003$ | $0.470 \pm 0.005$ |
| $g_1$ | $0.414 \pm 0.021$ | $0.735 \pm 0.034$ | $0.764 \pm 0.030$ | $0.701 \pm 0.031$ | $0.628 \pm 0.026$ | $1.250 \pm 0.056$ | $1.682 \pm 0.078$ |
| model II | EMM 28°C | EMM 30°C | EMM 32°C | EMM 34°C | YE 28°C | YE 30°C | YE 34°C |
| $\alpha$ | $2.068 \pm 0.152$ | $1.936 \pm 0.136$ | $2.068 \pm 0.146$ | $1.990 \pm 0.139$ | $1.419 \pm 0.082$ | $1.622 \pm 0.099$ | $1.518 \pm 0.090$ |
| $N$ | $16.387 \pm 0.800$ | $19.051 \pm 0.925$ | $18.609 \pm 0.898$ | $19.499 \pm 0.907$ | $30.137 \pm 1.382$ | $43.905 \pm 1.921$ | $50.067 \pm 2.037$ |
| $r_0$ | $0.635 \pm 0.010$ | $0.643 \pm 0.012$ | $0.644 \pm 0.014$ | $0.685 \pm 0.017$ | $0.702 \pm 0.020$ | $0.640 \pm 0.011$ | $0.614 \pm 0.009$ |
| $r_1$ | $0.089 \pm 0.008$ | $0.059 \pm 0.006$ | $0.054 \pm 0.006$ | $0.049 \pm 0.005$ | $0.072 \pm 0.007$ | $0.072 \pm 0.006$ | $0.068 \pm 0.007$ |
| $a$ | $0.004 \pm 0.001$ | $0.009 \pm 0.002$ | $0.006 \pm 0.001$ | $0.008 \pm 0.001$ | $0.117 \pm 0.017$ | $0.089 \pm 0.012$ | $0.151 \pm 0.022$ |
| $g_1$ | $0.380 \pm 0.017$ | $0.526 \pm 0.030$ | $0.710 \pm 0.037$ | $0.543 \pm 0.028$ | $0.444 \pm 0.017$ | $0.803 \pm 0.041$ | $0.936 \pm 0.046$ |
| $\nu$ | $225.97 \pm 8.68$ | $257.01 \pm 10.26$ | $201.98 \pm 7.56$ | $206.33 \pm 7.84$ | $198.97 \pm 6.99$ | $272.09 \pm 11.18$ | $270.18 \pm 10.85$ |

#### 1)

Estimation of *p* and *ν*. Note that the data of cell sizes just before division and just after division, *V_d_* and 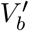, across different generations can be easily extracted from the lineage data and thus for model I, the parameter *p* can be estimated as the mean partition ratio 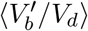. For model II, the parameters *p* and *ν* can be inferred by fitting the partition ratio data to a beta distribution. Typically, a mother cell divides into two daughters that are different in size due to stochasticity in partitioning and possible asymmetric cell division [31]. An interesting characteristic implied by the fission yeast data is that at division, the smaller daughter is always tracked with the mean partition ratio *p* being 0.46 – 0.47 for all the seven growth conditions (Table 2).

#### 2)

Estimation of *α*. Note that the data of cell sizes at birth and at division, *V_b_* and *V_d_*, across different generations can be easily extracted from the lineage data. For model I, since the parameter *p* has been determined, the strength *α* of cell size control can be estimated by finding the unique value of *α* satisfying the equality 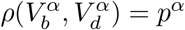. The inference of the control strength *α* for model II is much more complicated. Note that once *α* is determined, both *K*_1_ and *K*_2_ can be computed via Eq. (18). For model II, the mean and variance for the *α*th power of the birth size are given by (see Supplementary Section 4 for the proof)

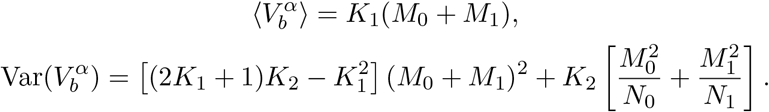

Since *K*_1_ and *K*_2_ have been determined (assuming *α* is known), it is possible to estimate both *M*_0_ + *M*_1_ and 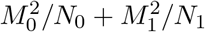 using the data of birth sizes. Finally, the control strength *α* can be estimated by finding the unique value of *α* satisfying Eq. (17).

#### 3)

Estimation of *g*_0_/*a* and *g*_1_/*a*. For model I, the mean and variance for the *α*th power of the birth size are given by (see Supplementary Section 4 for the proof)

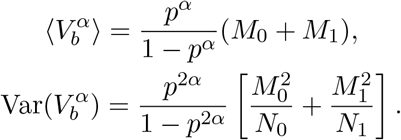

Since the parameters *p* and *α* have been determined, using the data of birth sizes, we are able to estimate the following two quantities:

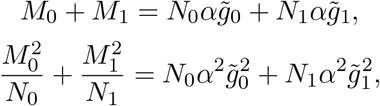

where 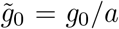 and 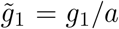. Once *N*_0_ and *N*_1_ are known, both 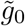 and 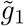 can be solved from the above two equations and thus can be inferred. For model II, we have shown how to estimate *M*_0_ + *M*_1_ and 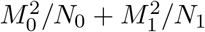 in step 2).

#### 4)

Estimation of *a, g*_0_, and *g*_1_. For each generation, say, the *k*th generation, we fit the time-course data of cell size to a three-stage growth model: an exponential growth in the elongation phase, followed by a constant size in the septation phase and another round of exponential growth in the fission phase:

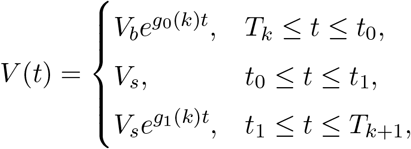

where *T_k_* and *T*_*k*+1_ are two successive division times, *g*_0_(*k*) and *g*_1_(*k*) are the growth rates in the elongation and fission phases for the *k*th generation, respectively, and *t*_0_ and *t*_1_ are the initial and end times of the septation phase, respectively. By carrying out least-squares optimal fitting, we can estimate the growth rate *g*_0_(*k*) in the elongation phase and the growth rate *g*_1_(*k*) in the fission phase for the *k*th generation. Fig. 5(a) illustrates the fitting of the time-course data to the three-stage growth model for three typical cell lineages, from which we can see that the model matches the data reasonably well. Then the parameter *g*_0_ can be determined as the mean of *g*_0_(*k*) across different generations. Since the time that the cell stays in the fission phase is very short, the estimate of *g*_1_(*k*) in general is not accurate. Therefore, we do not adopt this method to estimate the parameter *g*_1_. Since both *g*_0_ and 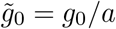 have been determined, the parameter a can also be inferred. Since both a and 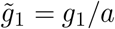 have been estimated, the parameter *g*_1_ can be determined.

#### 5)

Estimation of *N, N*_0_, and *N*_1_. Note that once the parameters *N, N*_0_, and *N*_1_ are known, all other parameters can be inferred by carrying out steps 1) - 4). Finally, we determine these three parameters by solving the following optimization problem:

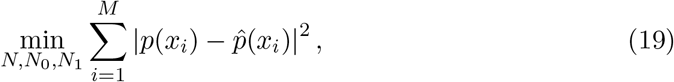

where *p*(*x*) is the theoretical cell size distribution, 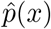 is the sample cell size distribution obtained from lineage data, *x_i_* are some reference points, and *M* is the number of bins chosen. In other words, we estimate the three parameters by matching the theoretical and experimental cell size distributions. For model I, the theoretical distribution is determined using Eq. (5), while for model II, the theoretical distribution is determined using Eq. (14). Thus far, all model parameters have been determined.

To test our parameter inference method, we compare the experimental cell size and birth size distributions obtained from lineage data (blue bars) with the theoretical ones based on the estimated parameters (red curves) under the seven growth conditions for both model I (Fig. 5(b),(c)) and model II (Fig. 6(a),(b)). It can be seen that the cell size distributions of lineage measurements for the seven growth conditions are all bimodal, while the birth size distributions are all unimodal. For the latter model, we also compare the distribution of the partition ratio with its approximation using the beta distribution (Fig. 6(c)). Clearly, the theory reproduces the experimental data of fission yeast excellently. Interestingly, while our inference method only involves the matching the theoretical and experimental cell size distribution, the theoretical birth size distribution also matches the experimental one reasonably well. The perfect match between theory and experiments supports the main assumptions of the three-stage growth model and the choice of the rate of moving from one cell cycle stage to the next to be a power law of cell size.

**Fig 5.**
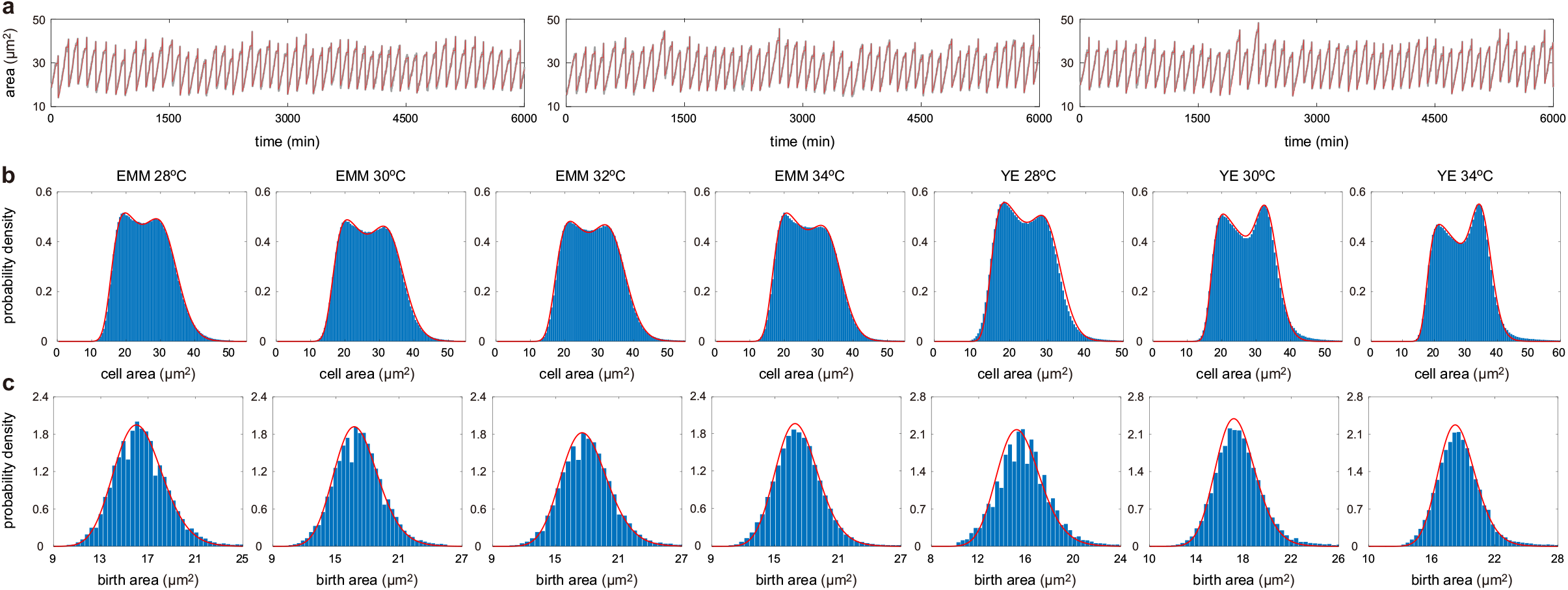
Fitting the experimental cell size and birth size distributions to theory based on the model with deterministic partitioning (model I). (**a**) Fitting the time-course data of cell size (grey curve) to a three-stage growth model (red curve) for three typical cell linages cultured in YE at 34°C. (**b**) Experimental cell size distributions (blue bars) and their optimal fitting to model I (red curve) for seven growth conditions. Here the theoretical distributions are computed using Eq. (5). (**c**) Same as (b) but for the birth size distributions. Here the theoretical distributions are computed using Eq. (12).

Our data analysis also reveals some significant differences between the two media used. From Table 2, it can be seen that cells cultured in EMM have a relatively strong size control (large *α*) and a relatively large added size variability (small *N*), while cells cultured in YE have a relatively weak size control (small *α*) and a relatively small added size variability (large *N*). Furthermore, we find that the size control strategy of fission yeast is sizer-like for all the seven growth conditions: for model II, the strength *α* of size control is typically 2.0 for EMM and is typically 1.5 for YE. This is in sharp contrast to the adder strategy found in *E. coli*, where *α* is estimated to be 0.8 – 1.2 for different growth conditions [13]. In addition, our data analysis predicts that the proportion of cell cycle stages in the elongation phase is about 60% – 70% and the proportion in the fission phase is about 5% – 10% for all growth conditions.

**Fig 6.**
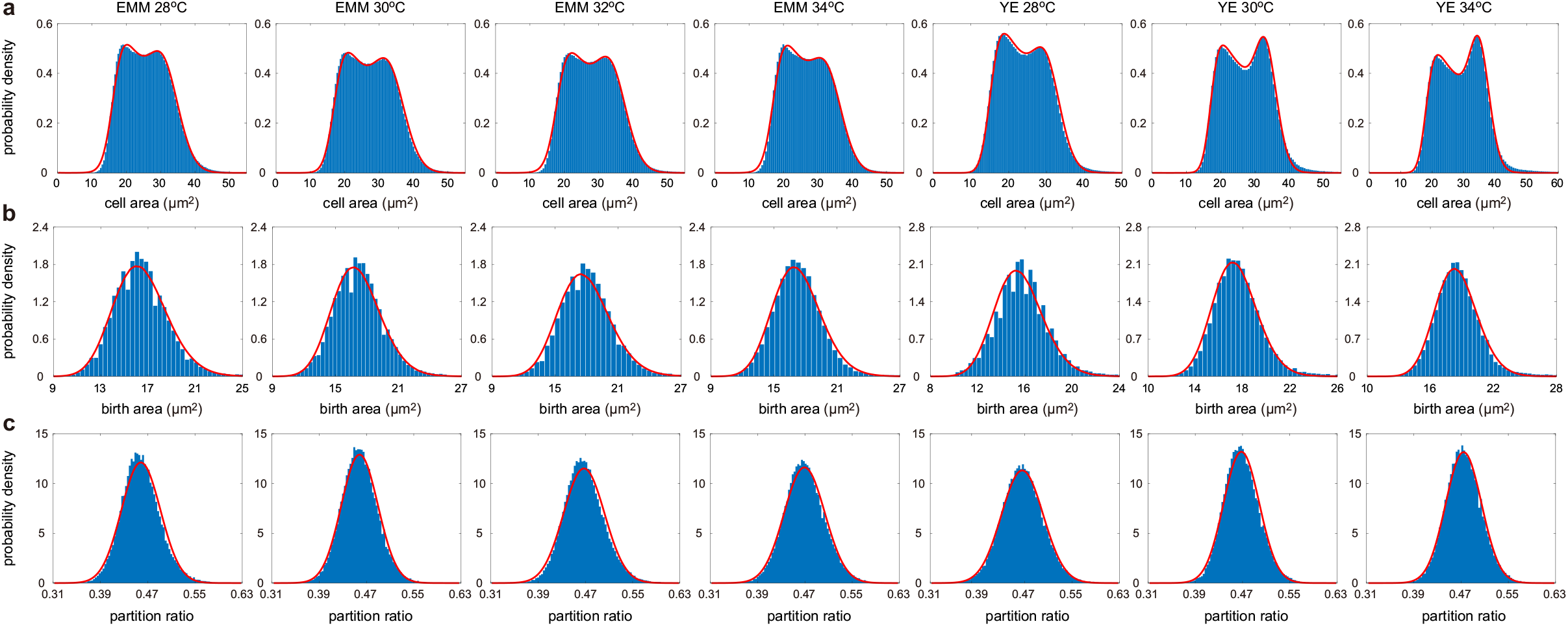
Fitting the experimental cell size, birth size, and partition ratio distributions to theory based on the model with stochastic partitioning (model II). (**a**) Experimental cell size distributions (blue bars) and their optimal fitting to model II (red curve) for seven growth conditions. Here the theoretical distributions are computed using Eq. (14). (**b**) Same as (b) but for the birth size distributions. Here the theoretical distributions are computed using stochastic simulations. (**c**) Same as (b) but for the partition ratio distributions. Here the theoretical distributions are computed using Eq. (3).

To further evaluate the performance of our model, we examine the correlation between cell sizes at birth and at division. Based on the lineage data, the correlation coefficients between birth and division sizes for the seven growth conditions are listed in the first row of Table 3. The theoretical predictions of the correlation coefficients based on stochastic simulations of model I and model II with the estimated parameters are listed in the second and third rows of Table 3, respectively. Clearly, both models capture the birth and division size correlations very well.

**Table 3.** Correlation coefficients between birth and division sizes for seven growth conditions. The experimental correlation coefficients are computed using the lineage data, while the theoretical correlation coefficients are computed using stochastic simulations based on model I and model II.

|  | EMM 28°C | EMM 30°C | EMM 32°C | EMM 34°C | YE 28°C | YE 30°C | YE 34°C |
| --- | --- | --- | --- | --- | --- | --- | --- |
| experiment | 0.2599 | 0.2834 | 0.2885 | 0.2753 | 0.4232 | 0.3534 | 0.3999 |
| model I | 0.2576 | 0.2704 | 0.2734 | 0.2844 | 0.4201 | 0.3544 | 0.3959 |
| model II | 0.2432 | 0.2604 | 0.2573 | 0.2740 | 0.4182 | 0.3669 | 0.4051 |

## Discussion

In this work, we have proposed two detailed models of cell size dynamics in fission yeast across many generations and analytically derived the cell size and birth size distributions of measurements obtained from a cell lineage. The main feature of cell size dynamics in fission yeast is its three-stage non-exponential growth pattern: a slow growth in the elongation phase, an arrest of growth in the septation phase, and a rapid growth in the fission phase. The first model assumes that (i) the cell undergoes deterministic exponential growth in the elongation and fission phases with the growth rate in the latter phase being greater than that in the former phase; (ii) the size remains constant in the septation phase; (iii) the size just after division is a fixed fraction of the one just before division; (iv) the cell cycle is divided into multiple effective cell cycle stages which correspond to different levels of the division protein (Cdc13, Cdc25, or Cdr2); (v) the rate of moving from one stage to the next has a power law dependence on cell size. A second model was also solved which relaxes assumption (iii) by allowing the size just after division to be a stochastic fraction of the one just before division with the fraction being distributed according to a beta distribution. Under assumptions (iv) and (v), the three typical strategies of size homeostasis (timer, adder, and sizer) are unified.

Experimentally, the cell size distribution of lineage data in fission yeast is typically bimodal under various growth conditions. This is very different from the unimodal size distribution obtained in many other cell types [13]. Interestingly, the bimodal cell size distribution of fission yeast can be excellently reproduced by the analytical solutions of both models. The origin of bimodality is further investigated and clarified in detail; we find that bimodality becomes apparent when (i) the variability in added size is not too large, (ii) the strength of size control is not too weak, which implies that adder or sizer-like strategies enforce size homeostasis, (iii) the proportion of the elongation phase in the cell cycle is neither too large nor too small, (iv) the proportion of the septation phase is large, (v) the proportion of the fission phase is small, and (vi) the size addition in the fission phase is not too sharp. We also find that fluctuations in partitioning at division has a considerable influence on the shape of the cell size distribution by declining the slope of the left shoulder, as well as lowering the heights of the two peaks.

Furthermore, we have developed an effective method of inferring all the parameters involved in both models using single-cell lineage measurements of fission yeast based on the information of (i) the partition ratio, namely, the ratio of the size just after division to the size just before division, across different generations, (ii) the mean and variance of the birth size across different generations, (iii) the correlation of cell sizes at birth and at division, and (iv) the cell size distribution. Specifically, we infer the parameters except the numbers of cell cycle stages in different phases using the information (i)-(iii) and then determine the remaining parameters by matching the theoretical and experimental cell size distributions.

We have shown that the theoretical cell size and birth size distributions provide an excellent fit to the experimental ones of fission yeast reported in [11] under seven different growth conditions. This match provides support for two implicit important assumptions of our model: (i) the cell undergoes a complex three-stage growth pattern and (ii) the speed of the cell cycle progression (the transition rate between cell cycle stages) has a power law dependence on cell size. Finally, based on matching the experimental to the theoretical cell size distributions, we have estimated all model parameters from lineage data of fission yeast and found that the variability in added size and the strength of size control are remarkably different when cells are cultured in different media — EMM has a large added size variability and a strong size control, while YE has a small added size variability and a weak size control. The estimated values of the strength *α* of size homeostasis is typically 2.0 for EMM and 1.5 for YE, confirming the previous results that fission yeast uses the sizer-like strategy to achieve size homeostasis [16]. Simulations with the inferred parameters using distribution matching also captured the correlation between birth and division sizes — this provides further evidence of the accuracy of our detailed model.

## Data Availability

All data needed to evaluate the conclusions in the paper are present in the paper and in [11].

## Author contributions

R. G. conceived the original idea. C. J. performed the theoretical derivations and numerical simulations. C. J, A. S, and R. G interpreted the theoretical results. C. J and R. G jointly wrote the manuscript with input from A. S.

## Acknowledgments

C. J. acknowledges support from the NSAF grant in National Natural Science Foundation of China with grant No. U1930402. A. S. is supported by the National Institute of Health Grant 1R01GM126557. R. G. acknowledges support from the Leverhulme Trust (RPG-2018-423).

