## Supplemental Information for "Characterizing non-exponential growth and bimodal cell size distributions in Schizosaccharomyces pombe: an analytical approach"

#### 1 Distribution of the generalized added size

Let $V_b$ and $V_d$ denote cell sizes at birth and at division in a particular generation, respectively, and let $V_s$ denote cell size in the septation phase, which is assumed to be a constant. In the main text, we have stated that (i) the generalized added size $V_s^\alpha - V_b^\alpha$ in the elongation phase is Erlang distributed with shape parameter $N_0$ and mean $M_0 = N_0 g_0 \alpha/a$ and (ii) the generalized added size $V_d^\alpha - V_s^\alpha$ in the fission phase is also Erlang distributed with shape parameter $N_1$ and mean $M_1 = N_1 g_1 \alpha/a$. To see this, note that when $V_b$ is fixed, the cell size in the elongation phase is given by $V(t) = V_b e^{g_0 t}$. Since the transition rate from one stage to the next at time $t$ is equal to $aV(t)^\alpha$, the distribution of the transition time $T$ is given by

$$\mathbb{P}(T > t) = e^{-\int_0^t aV(s)^\alpha ds} = e^{-\int_0^t aV_b^\alpha e^{g_0 \alpha s} ds} = e^{-\frac{aV_b^\alpha}{g_0 \alpha}(e^{g_0 \alpha t}-1)}. \tag{1}$$

This shows that

$$\mathbb{P}(V_b^\alpha(e^{g_0 \alpha T} - 1) > t) = e^{-\frac{at}{g_0 \alpha}}.$$

Hence $V_b^\alpha(e^{g_0 \alpha T} - 1)$ is exponentially distributed with mean $g_0 \alpha/a$. Note that $V_b^\alpha(e^{g_0 \alpha T} - 1)$ is the generalized added size in a particular cell cycle stage. As a result, the generalized added size in the elongation phase, $V_s^\alpha - V_b^\alpha$, is the independent sum of $N_0$ exponentially distributed random variables with mean $g_0 \alpha/a$. This shows that $V_s^\alpha - V_b^\alpha$ has an Erlang distribution with shape parameter $N_0$ and mean $M_0 = N_0 g_0 \alpha/a$. The fact that $V_d^\alpha - V_s^\alpha$ has an Erlang distribution with shape parameter $N_1$ and mean $M_1 = N_1 g_1 \alpha/a$ can be proved in the same way.

#### 2 Cell size distribution

##### 2.1 Model with deterministic partitioning: general case

Here we compute the cell size distribution of lineage measurements for the model with deterministic partitioning. The microstate of the cell can be represented by an ordered pair $(k, x)$, where $k$ is the cell cycle stage and $x$ is the cell size. Let $p_k(x)$ denote the probability density function of cell size when the cell is in stage $k$. Then the evolution of cell size dynamics is governed by the master equation

$$\begin{aligned}
&\partial_t p_1(x) = -\partial_x[g_0 x p_1(x)] + \frac{a}{p}\left(\frac{x}{p}\right)^{\alpha} p_N\left(\frac{x}{p}\right) - a x^{\alpha} p_1(x),\\
&\partial_t p_k(x) = -\partial_x[g_0 x p_k(x)] + a x^{\alpha} p_{k-1}(x) - a x^{\alpha} p_k(x), \quad 2 \le k \le N_0,\\
&\partial_t p_k(x) = a x^{\alpha} p_{k-1}(x) - a x^{\alpha} p_k(x), \quad N_0 + 1 \le k \le N - N_1,\\
&\partial_t p_k(x) = -\partial_x[g_1 x p_k(x)] + a x^{\alpha} p_{k-1}(x) - a x^{\alpha} p_k(x), \quad N - N_1 + 1 \le k \le N.
\end{aligned}$$

For the first, second, and fourth equations, the first term on the right-hand side describe cell growth and the remaining two terms describe transitions between cell cycle stages. For the third equation, the two terms on the right-hand side describe cell cycle stage transitions. For the first equation, the middle term on the right-hand side describes partitioning of cell size at division.

For convenience, we next focus on the $\alpha$th power of cell size, $y = x^{\alpha}$. Let $\tilde{p}_k(y)$ denote the probability density function of the $\alpha$th power of cell size when the cell is in stage $k$. It is easy to see that $p_k(x)$ and $\tilde{p}_k(y)$ are related by

$$\tilde{p}_k(y) = \frac{1}{\alpha} y^{\frac{1}{\alpha}-1} p_k(y^{\frac{1}{\alpha}}).$$

In other words, we have

$$\alpha y \tilde{p}_k(y) = x p_k(x).$$

Based on this relation, the master equation can be rewritten as

$$\begin{aligned}
&\partial_t \tilde{p}_1(y) = -\partial_y[\alpha g_0 y \tilde{p}_1(y)] + \frac{ay}{p^{2\alpha}} \tilde{p}_N\left(\frac{y}{p^{\alpha}}\right) - ay\tilde{p}_1(y),\\
&\partial_t \tilde{p}_k(y) = -\partial_y[\alpha g_0 y \tilde{p}_k(y)] + ay\tilde{p}_{k-1}(y) - ay\tilde{p}_k(y), \quad 2 \le k \le N_0,\\
&\partial_t \tilde{p}_k(y) = ay\tilde{p}_{k-1}(y) - ay\tilde{p}_k(y), \quad N_0 + 1 \le k \le N - N_1,\\
&\partial_t \tilde{p}_k(y) = -\partial_y[\alpha g_1 y \tilde{p}_k(y)] + ay\tilde{p}_{k-1}(y) - ay\tilde{p}_k(y), \quad N - N_1 + 1 \le k \le N.
\end{aligned} \tag{2}$$

To proceed, for each cell cycle stage $k$, we introduce the Laplace transform

$$\begin{aligned}
F_k(\lambda) &= \int_0^{\infty} \tilde{p}_k(y) e^{-\lambda y} dy = \int_0^{\infty} p_k(x) e^{-\lambda x^{\alpha}} dx,\\
F(\lambda) &= \int_0^{\infty} \tilde{p}(y) e^{-\lambda y} dy = \int_0^{\infty} p(x) e^{-\lambda x^{\alpha}} dx,
\end{aligned} \tag{3}$$

where $\tilde{p}(y) = \sum_{k=1}^{N} \tilde{p}_k(y)$. Then Eq. (2) can be converted to the following differential equations:

$$\begin{aligned}
&\partial_t F_1(\lambda) = (\alpha g_0 \lambda + a)\partial_\lambda F_1(\lambda) - a\partial_\lambda F_N(p^{\alpha}\lambda),\\
&\partial_t F_k(\lambda) = (\alpha g_0 \lambda + a)\partial_\lambda F_k(\lambda) - a\partial_\lambda F_{k-1}(\lambda), \quad 2 \le k \le N_0,\\
&\partial_t F_k(\lambda) = a\partial_\lambda F_k(\lambda) - a\partial_\lambda F_{k-1}(\lambda), \quad N_0 + 1 \le k \le N - N_1,\\
&\partial_t F_k(\lambda) = (\alpha g_1 \lambda + a)\partial_\lambda F_k(\lambda) - a\partial_\lambda F_{k-1}(\lambda), \quad N - N_1 + 1 \le k \le N.
\end{aligned}$$

At the steady state, the above equations reduce to

$$\begin{aligned}
&F'_N(p^{\alpha}\lambda) = (1 + A_0\lambda)F'_1(\lambda),\\
&F'_{k-1}(\lambda) = (1 + A_0\lambda)F'_k(\lambda), \quad 2 \le k \le N_0,\\
&F'_{k-1}(\lambda) = F'_k(\lambda), \quad N_0 + 1 \le k \le N - N_1,\\
&F'_{k-1}(\lambda) = (1 + A_1\lambda)F'_k(\lambda), \quad N - N_1 + 1 \le k \le N,
\end{aligned} \tag{4}$$

where $A_0 = \alpha g_0/a$ and $A_1 = \alpha g_1/a$. Note the the above equations are recursive with respect to $k$, which means that $F'_{k-1}$ can be represented by $F'_k$ for each $2 \leq k \leq N$. Using the recursive relations repeatedly, we find that $F'_k$ can be represented by $F'_N$ as

$$\begin{aligned} F'_k(\lambda) &= (1+A_1\lambda)^{N-k} F'_N(\lambda), \quad N-N_1 \leq k \leq N, \\ F'_k(\lambda) &= (1+A_1\lambda)^{N_1} F'_N(\lambda), \quad N_0 \leq k \leq N-N_1-1, \\ F'_k(\lambda) &= (1+A_0\lambda)^{N_0-k}(1+A_1\lambda)^{N_1} F'_N(\lambda), \quad 1 \leq k \leq N_0-1. \end{aligned} \tag{5}$$

In particular, $F'_1$ can be represented by $F'_N$ as

$$F'_1(\lambda) = (1+A_0\lambda)^{N_0-1}(1+A_1\lambda)^{N_1} F'_N(\lambda).$$

Inserting this equation into the first equality of Eq. (4), we obtain

$$F'_N(\lambda) = b(\lambda) F'_N(p^\alpha \lambda), \tag{6}$$

where

$$b(\lambda) = (1+A_0\lambda)^{-N_0}(1+A_1\lambda)^{-N_1}. \tag{7}$$

This is a functional equation satisfied by $F_N$. Applying Eq. (6) repeatedly, we obtain

$$F'_N(\lambda) = \prod_{k=0}^{n-1} b(p^{\alpha k}\lambda) F'_N(p^{\alpha n}\lambda), \quad n \geq 1.$$

Taking $n \to \infty$ in the above equation yields

$$F'_N(\lambda) = F'_N(0) \prod_{k=0}^{\infty} b(p^{\alpha k}\lambda). \tag{8}$$

On the other hand, summing over all the equalities in Eq. (5), we obtain

$$F'(\lambda) = N f(\lambda) F'_N(\lambda), \tag{9}$$

where

$$f(\lambda) = (1+A_1\lambda)^{N_1}\left[\frac{(1+A_0\lambda)^{N_0}-1}{NA_0\lambda} + \frac{N-N_0-N_1}{N}\right] + \frac{(1+A_1\lambda)^{N_1}-1}{NA_1\lambda}. \tag{10}$$

Combining Eqs. (8) and (9), we obtain

$$F'(\lambda) = NF'_N(0) f(\lambda) \prod_{k=0}^{\infty} b(p^{\alpha k}\lambda).$$

Since $F(\infty) = 0$, integrating the above equation yields

$$F(\lambda) = -NF'_N(0) \int_\lambda^\infty f(u) \prod_{k=0}^{\infty} b(p^{\alpha k}u) du.$$

Finally, using the fact that $F(0) = 1$, we obtain the explicit expression of the Laplace transform, which is given by

$$F(\lambda) = K \int_\lambda^\infty f(u) \prod_{k=0}^{\infty} b(p^{\alpha k}u) du. \tag{11}$$

where

$$K = -NF_N'(0) = \left[\int_0^\infty f(u) \prod_{k=0}^\infty b(p^{\alpha k}u)du\right]^{-1}.$$

is a normalization constant. Taking the inverse Laplace transform of $F(\lambda)$ gives the probability density function $\tilde{p}(y)$ of the $\alpha$th power of cell size. Finally, the probability density function $p(x)$ of the original cell size is given by

$$p(x) = \alpha x^{\alpha-1}\tilde{p}(x^\alpha). \tag{12}$$

Next we focus on how to compute the cell size distribution numerically. Taking the derivative with respect to $\lambda$ on both sides of Eq. (11) yields

$$\int_0^\infty y\tilde{p}(y)e^{-\lambda y}dy = Kf(\lambda)\prod_{k=0}^\infty b(p^{\alpha k}\lambda) := G(\lambda).$$

Replacing $\lambda$ by $i\lambda$ in the above equation yields

$$\int_0^\infty y\tilde{p}(y)e^{-i\lambda y}dy = Kf(i\lambda)\prod_{k=0}^\infty b(p^{\alpha k}i\lambda) := G(i\lambda).$$

This shows that the Fourier transform of $y\tilde{p}(y)$ is exactly $G(i\lambda)$. Since the Fourier transform and inverse fourier transform are inverses of each other, we only need to take the inverse Fourier transform of $G(i\lambda)$ to obtain $y\tilde{p}(y)$. Finally, we use Eq. (12) to obtain the cell size distribution $p(x)$.

#### 2.2 Model with deterministic partitioning: limiting case

We next focus on the special case where the cell cycle variability is very small ($N \gg 1$). In this limit, the function $b(\lambda)$ reduces to

$$b(\lambda) = \left(\frac{M_0\lambda}{N_0}+1\right)^{-N_0}\left(\frac{M_1\lambda}{N_1}+1\right)^{-N_1} = e^{-(M_0+M_1)\lambda},$$

and the function $f(\lambda)$ reduces to

$$f(\lambda) = e^{M_1\lambda}\left[\frac{r_0(e^{M_0\lambda}-1)}{M_0\lambda}+1-r_0-r_1\right]+\frac{r_1(e^{M_1\lambda}-1)}{M_1\lambda}, \tag{13}$$

where $r_0 = N_0/N$ is the proportion of the elongation phase and $r_1 = N_1/N$ is the proportion of the fission phase. This shows that

$$\prod_{k=0}^\infty b(p^{\alpha k}\lambda) = \prod_{k=0}^\infty e^{-(M_0+M_1)p^{\alpha k}\lambda} = e^{-\frac{(M_0+M_1)\lambda}{1-p^\alpha}}. \tag{14}$$

Combining Eqs. (13) and (14), it is easy to check that

$$f(\lambda)\prod_{k=0}^\infty b(p^{\alpha k}\lambda) = \frac{r_0(e^{-v_b^\alpha u}-e^{-v_m^\alpha u})}{M_0u}+(1-r_0-r_1)e^{-v_m^\alpha u}+\frac{r_1(e^{-v_b^\alpha u}-e^{-v_m^\alpha u})}{M_1u}.$$

where $v_b$, $v_m$, and $v_d$ are constants defined as

$$v_b = p\left(\frac{M_0+M_1}{1-p^\alpha}\right)^{\frac{1}{\alpha}},\quad v_m = \left(\frac{M_0+M_1p^\alpha}{1-p^\alpha}\right)^{\frac{1}{\alpha}},\quad v_d = \left(\frac{M_0+M_1}{1-p^\alpha}\right)^{\frac{1}{\alpha}}.$$

Thus the Laplace transform $F(\lambda)$ can be simplified as

$$F(\lambda) = K\int_\lambda^\infty \left[\frac{r_0(e^{-v_b^\alpha u}-e^{-v_m^\alpha u})}{M_0u}+(1-r_0-r_1)e^{-v_m^\alpha u}+\frac{r_1(e^{-v_b^\alpha u}-e^{-v_m^\alpha u})}{M_1u}\right]du$$
$$= \frac{Kr_0}{M_0}[E_1(v_b^\alpha\lambda)-E_1(v_m^\alpha\lambda)]+\frac{K(1-r_0-r_1)}{v_m^\alpha}e^{-v_m^\alpha\lambda}+\frac{Kr_1}{M_1}[E_1(v_m^\alpha\lambda)-E_1(v_d^\alpha\lambda)],$$

where

$$E_1(x) = \int_x^\infty \frac{e^{-u}}{u}du$$

is the exponential integral and

$$K = \left\{\int_0^\infty \left[\frac{r_0(e^{-v_b^\alpha u}-e^{-v_m^\alpha u})}{M_0u}+(1-r_0-r_1)e^{-v_m^\alpha u}+\frac{r_1(e^{-v_b^\alpha u}-e^{-v_m^\alpha u})}{M_1u}\right]du\right\}^{-1}$$

is a normalization constant. The normalization constant can be calculated explicitly as

$$K = (T_1+T_2+T_3)^{-1},$$

where $T_1$, $T_2$, and $T_3$ are constants defined as

$$T_1 = \frac{\alpha r_0}{M_0}\log\frac{v_m}{v_b},\quad T_2 = \frac{1-r_0-r_1}{v_m^\alpha},\quad T_3 = \frac{\alpha r_1}{M_1}\log\frac{v_d}{v_m}.$$

Taking the inverse Laplace transform finally gives the distribution of the $\alpha$th power of cell size

$$\tilde{p}(y) = \frac{w_1}{\alpha(\log v_m-\log v_b)y}I_{[v_b^\alpha,v_m^\alpha]}(y)+w_2\delta(y-v_m^\alpha)+\frac{w_3}{\alpha(\log v_d-\log v_m)y}I_{[v_m^\alpha,v_d^\alpha]}(y),$$

where $I_A(x)$ is the indicator function which takes the value of 1 when $x \in A$ and takes the value of 0 otherwise, $\delta(x)$ is Dirac's delta function, and $w_1$, $w_2$, and $w_3$ are constants defined as

$$w_1 = \frac{T_1}{T_1+T_2+T_3},\quad w_2 = \frac{T_2}{T_1+T_2+T_3},\quad w_3 = \frac{T_3}{T_1+T_2+T_3}.$$

Finally, the distribution of cell size is given by

$$p(x) = \frac{w_1}{(\log v_m-\log v_b)x}I_{[v_b,v_m]}(x)+w_2\delta(x-v_m)+\frac{w_3}{(\log v_d-\log v_m)x}I_{[v_m,v_d]}(x), \tag{15}$$

where $w_1$, $w_2$, and $w_3$ represent the proportions of subpopulations in the elongation, septation, and fission phases, respectively and $v_b$, $v_m$, and $v_d$ represent the typical birth, septation, and division sizes, respectively. The analytical distribution can be used to compute some other quantities of interest. For example, when $N \gg 1$, the mean cell size is given by

$$\langle V\rangle = \frac{w_1(v_m-v_b)}{\log v_m-\log v_b}+w_2v_m+\frac{w_3(v_d-v_m)}{\log v_d-\log v_m}.$$

#### 2.3 Model with stochastic partitioning

Next we compute the cell size distribution of lineage measurements for the model with stochastic partitioning. In this case, the evolution of cell size dynamics is governed by the master equation

$$\partial_t p_1(x) = -\partial_x[g_0xp_1(x)]+\int_0^1 \frac{a}{z}\left(\frac{x}{z}\right)^\alpha p_N\left(\frac{x}{z}\right)f(z)dz-ax^\alpha p_1(x),$$
$$\partial_t p_k(x) = -\partial_x[g_0xp_k(x)]+ax^\alpha p_{k-1}(x)-ax^\alpha p_k(x),\quad 2\le k\le N_0,$$
$$\partial_t p_k(x) = ax^\alpha p_{k-1}(x)-ax^\alpha p_k(x),\quad N_0+1\le k\le N-N_1,$$
$$\partial_t p_k(x) = -\partial_x[g_1xp_k(x)]+ax^\alpha p_{k-1}(x)-ax^\alpha p_k(x),\quad N-N_1+1\le k\le N.$$

For convenience, we next focus on the $\alpha$th power of cell size, $y = x^\alpha$. Let $\tilde{p}_k(y)$ denote the probability density function of the $\alpha$th power of cell size when the cell is in stage $k$. Then the master equation can be rewritten as

$$\begin{aligned}
&\partial_t\tilde{p}_1(y) = -\partial_y[\alpha g_0 y\tilde{p}_1(y)] + \int_0^1 \frac{ay}{z^{2\alpha}}\tilde{z}_N\left(\frac{y}{z^\alpha}\right)f(z)dz - ay\tilde{p}_1(y),\\
&\partial_t\tilde{p}_k(y) = -\partial_y[\alpha g_0 y\tilde{p}_k(y)] + ay\tilde{p}_{k-1}(y) - ay\tilde{p}_k(y), \quad 2 \le k \le N_0,\\
&\partial_t\tilde{p}_k(y) = ay\tilde{p}_{k-1}(y) - ay\tilde{p}_k(y), \quad N_0+1 \le k \le N-N_1,\\
&\partial_t\tilde{p}_k(y) = -\partial_y[\alpha g_1 y\tilde{p}_k(y)] + ay\tilde{p}_{k-1}(y) - ay\tilde{p}_k(y), \quad N-N_1+1 \le k \le N.
\end{aligned}$$

Using the Laplace transform defined in Eq. (3), the above equations can be converted to the following differential equations:

$$\begin{aligned}
&\partial_t F_1(\lambda) = (\alpha g_0\lambda + a)\partial_\lambda F_1(\lambda) - a\int_0^1 \partial_\lambda F_N(z^\alpha\lambda)f(z)dz,\\
&\partial_t F_k(\lambda) = (\alpha g_0\lambda + a)\partial_\lambda F_k(\lambda) - a\partial_\lambda F_{k-1}(\lambda), \quad 2 \le k \le N_0,\\
&\partial_t F_k(\lambda) = a\partial_\lambda F_k(\lambda) - a\partial_\lambda F_{k-1}(\lambda), \quad N_0+1 \le k \le N-N_1,\\
&\partial_t F_k(\lambda) = (\alpha g_1\lambda + a)\partial_\lambda F_k(\lambda) - a\partial_\lambda F_{k-1}(\lambda), \quad N-N_1+1 \le k \le N.
\end{aligned}$$

At the steady state, the above equations reduce to

$$\begin{aligned}
&\int_0^1 F_N'(z^\alpha\lambda)dz = (1 + A_0\lambda)F_1'(\lambda),\\
&F_{k-1}'(\lambda) = (1 + A_0\lambda)F_k'(\lambda), \quad 2 \le k \le N_0,\\
&F_{k-1}'(\lambda) = F_k'(\lambda), \quad N_0+1 \le k \le N-N_1,\\
&F_{k-1}'(\lambda) = (1 + A_1\lambda)F_k'(\lambda), \quad N-N_1+1 \le k \le N,
\end{aligned}$$

where $A_0 = \alpha g_0/a$ and $A_1 = \alpha g_1/a$. In analogy to the derivation for model I, we have

$$F_N'(\lambda) = b(\lambda)\int_0^1 F_N'(z^\alpha\lambda)dz, \tag{16}$$

where $b(\lambda)$ is the function given in Eq. (7). This is a functional integral equation satisfied by $F_N$. To solve this, we expand both $F_N'(\lambda)$ and $b(\lambda)$ in power series as

$$F_N'(\lambda) = \sum_{n=0}^\infty x_n\lambda^n, \quad b(\lambda) = \sum_{n=0}^\infty b_n\lambda^n. \tag{17}$$

Using the functional form of $b(\lambda)$ in Eq. (7), it is not hard to check that

$$b_n = \frac{(-1)^n}{n!}\sum_{m=0}^n C_{n,m}(N_0)_m(N_1)_{n-m}A_0^m A_1^{n-m}, \tag{18}$$

where $C_{n,m} = n!/m!(n-m)!$ is the combinatorial number and $(x)_m = x(x+1)\cdots(x+m-1)$ is the Pochhammer symbol. Inserting Eq. (17) into Eq. (16), we obtain

$$\sum_{n=0}^\infty x_n\lambda^n = \sum_{n=0}^\infty b_n\lambda^n\sum_{n=0}^\infty x_n c_n\lambda^n = \sum_{n=0}^\infty\sum_{m=0}^n x_m c_m b_{n-m}\lambda^n, \tag{19}$$

where

$$c_n = \int_0^1 z^{\alpha n} f(z) dz = \frac{B(\alpha n + p\nu, q\nu)}{B(p\nu, q\nu)}. \tag{20}$$

Comparing the coefficients on both sides of Eq. (19), we obtain

$$x_n = \sum_{m=0}^{n} x_m c_m b_{n-m}.$$

This can be rewritten as

$$x_n = F_N'(0) a_n,$$

where $a_n$ can be determined using the following recursive relations:

$$a_n = \frac{1}{1-c_n} \sum_{m=0}^{n-1} a_m c_m b_{n-m}, \quad a_0 = 1, \tag{21}$$

where $b_n$ is the sequence defined in Eq. (18) and $c_n$ is the sequence defined in Eq. (20). To summarize, we obtain

$$F_N'(\lambda) = F_N'(0) \sum_{n=0}^{\infty} a_n \lambda^n, \tag{22}$$

where $a_n$ is the sequence defined in Eq. (21).

On the other hand, in analogy to the derivation for model I, we have

$$F'(\lambda) = N f(\lambda) F_N'(\lambda), \tag{23}$$

where $f(\lambda)$ is the function given in Eq. (10). Combining Eqs. (22) and (23), we obtain

$$F'(\lambda) = N F_N'(0) f(\lambda) \sum_{n=0}^{\infty} a_n \lambda^n.$$

Since $F(\infty) = 0$, integrating the above equation yields

$$F(\lambda) = -N F_N'(0) \int_{\lambda}^{\infty} f(u) \sum_{n=0}^{\infty} a_n u^n du.$$

Finally, using the fact that $F(0) = 1$, we obtain the explicit expression of the Laplace transform, which is given by

$$F(\lambda) = K \int_{\lambda}^{\infty} f(u) \sum_{n=0}^{\infty} a_n u^n du. \tag{24}$$

where

$$K = -N F_N'(0) = \left[ \int_0^{\infty} f(u) \sum_{n=0}^{\infty} a_n u^n du \right]^{-1}.$$

is a normalization constant. Taking the inverse Laplace transform of $F(\lambda)$ gives the probability density function $\tilde{p}(y)$ of the $\alpha$th power of cell size. Finally, the probability density function $p(x)$ of the original cell size is given by

$$p(x) = \alpha x^{\alpha-1} \tilde{p}(x^\alpha).$$

For the special case of exponential growth of cell size, there is only the elongation phase and the septation and fission phases vanish. In this case, we have $N_1 = 0$ and $N = N_0$; the cell size distribution is still determined by Eq. (24) with the sequence $b_n$ and the function $f(\lambda)$ being simplified as

$$b_n = \frac{(N)_n (-A_0)^n}{n!}, \quad f(\lambda) = \frac{(1 + A_0\lambda)^N - 1}{N A_0 \lambda}.$$

### 3 Birth size distribution

Here we compute the distribution of $V_b$. To this end, let $V_b(k)$ and $V_d(k)$ denote the cell sizes at birth and at division in the $k$th generation, respectively. Under the assumption of deterministic partitioning, we have $V_b(k+1) = pV_d(k)$ and thus we obtain the recursive equation

$$V_b^\alpha(k+1) = p^\alpha[V_b^\alpha(k) + \Delta_k], \quad k \geq 0,$$

where $\Delta_k = V_d^\alpha(k) - V_b^\alpha(k)$ is the generalized added size in the $k$th generation. Using the recursive equation repeatedly, we obtain

$$V_b^\alpha(k) = p^{k\alpha}V_b(0)^\alpha + p^{k\alpha}\Delta_0 + p^{(k-1)\alpha}\Delta_1 + \cdots + p^\alpha\Delta_{k-1}. \tag{25}$$

Recall that $\Delta_0, \Delta_1, \Delta_2, \cdots$ are i.i.d. hypoexponentially distributed random variables with the Laplace transform of each term being given by

$$\mathbb{E}e^{-\lambda\Delta_n} = b(\lambda).$$

It thus follows from (25) and the independence of $V_b(0), \Delta_0, \Delta_1, \Delta_2, \cdots$ that

$$\mathbb{E}e^{-\lambda V_b^\alpha(k)} = \mathbb{E}e^{-\lambda p^{k\alpha}V_b(0)^\alpha}\prod_{n=1}^{k} b(p^{n\alpha}\lambda). \tag{26}$$

Since the distribution of $V_b(k)$ converges to the steady-state distribution of the birth size as $k \to \infty$, taking $k \to \infty$ in Eq. (26) shows that the Laplace transform of $V_b^\alpha$ is given by

$$\mathbb{E}e^{-\lambda V_b^\alpha} = \prod_{n=1}^{\infty} b(p^{\alpha n}u) = \prod_{n=1}^{\infty}\left(1 + \frac{M_0p^{\alpha n}\lambda}{N_0}\right)^{-N_0}\left(1 + \frac{M_1p^{\alpha n}\lambda}{N_1}\right)^{-N_1}. \tag{27}$$

Taking the inverse Laplace transform gives the probability density function of $V_b^\alpha$, from which is the probability density function of $V_b$ can be obtained.

### 4 Correlation between birth and division sizes

#### 4.1 Model with deterministic partitioning

Let $V_b$ and $V_d$ denote the cell sizes at birth and at division in a particular generation, respectively, and let $V_b'$ and $V_d'$ denote the the birth and division sizes in the next generation, respectively. We first focus on the correlation between the birth size $V_d$ and the division size $V_d$ for the model with deterministic partitioning (model I). Since the generalized added size $\Delta = V_d^\alpha - V_b^\alpha$ is independent of $V_b$, we have

$$\mathrm{Cov}(V_b^\alpha, V_d^\alpha) = \mathrm{Cov}(V_b^\alpha, V_b^\alpha + \Delta) = \mathrm{Var}(V_b^\alpha),$$

as well as

$$\mathrm{Var}(V_d^\alpha) = \mathrm{Var}(V_b^\alpha + \Delta) = \mathrm{Var}(V_b^\alpha) + \mathrm{Var}(\Delta),$$

where $\mathrm{Cov}(X, Y)$ denotes the covariance between random variables $X$ and $Y$ and $\mathrm{Var}(X)$ denotes the variance of $X$. This shows that

$$\rho(V_b^\alpha, V_d^\alpha) = \frac{\mathrm{Cov}(V_b^\alpha, V_d^\alpha)}{\sqrt{\mathrm{Var}(V_b^\alpha)\mathrm{Var}(V_d^\alpha)}} = \sqrt{\frac{\mathrm{Var}(V_b^\alpha)}{\mathrm{Var}(V_b^\alpha) + \mathrm{Var}(\Delta)}}, \tag{28}$$

where $\rho(X,Y)$ denotes the covariance between $X$ and $Y$. Since the generalized added size $\Delta$ is the independent sum of an Erlang distributed random variable with shape parameter $N_0$ and mean $M_0$ and another Erlang distributed random variable with shape parameter $N_1$ and mean $M_1$, we have

$$\mathrm{Var}(\Delta) = \frac{M_0^2}{N_0} + \frac{M_1^2}{N_1}. \tag{29}$$

Moreover, since $V_b' = pV_d$, we have

$$V_b'^{\alpha} = p^{\alpha}(V_b^{\alpha} + \Delta).$$

This shows that $p^{\alpha}(V_b^{\alpha} + \Delta)$ and $V_b^{\alpha}$ have the same distribution. Thus we obtain

$$\mathbb{E}V_b^{\alpha} = p^{\alpha}\mathbb{E}(V_b^{\alpha} + \Delta), \tag{30}$$

$$\mathbb{E}V_b^{2\alpha} = p^{2\alpha}\mathbb{E}(V_b^{\alpha} + \Delta)^2. \tag{31}$$

It then follows from Eq. (35) that

$$\mathbb{E}V_b^{\alpha} = \frac{p^{\alpha}}{1-p^{\alpha}}(M_0 + M_1). \tag{32}$$

Similarly, it follows from Eq. (36) that

$$\mathbb{E}V_b^{2\alpha} = \frac{p^{2\alpha}}{(1-p^{\alpha})^2}(M_0 + M_1)^2 + \frac{p^{2\alpha}}{1-p^{2\alpha}}\left[\frac{M_0^2}{N_0} + \frac{M_1^2}{N_1}\right]. \tag{33}$$

Combining Eqs. (37) and (38) shows that

$$\mathrm{Var}(V_b^{\alpha}) = \mathbb{E}V_b^{2\alpha} - (\mathbb{E}V_b^{\alpha})^2 = \frac{p^{2\alpha}}{1-p^{2\alpha}}\left[\frac{M_0^2}{N_0} + \frac{M_1^2}{N_1}\right].$$

Inserting this equation into Eq. (28) finally shows that

$$\rho(V_b^{\alpha}, V_d^{\alpha}) = \sqrt{p^{2\alpha}} = p^{\alpha}. \tag{34}$$

We next focus on the correlation between two successive birth sizes and the correlation between two successive division sizes. Since $V_b' = pV_d$, the correlation coefficient between $V_b^{\alpha}$ and $V_b'^{\alpha}$ is exactly the same as that between $V_b^{\alpha}$ and $V_d^{\alpha}$, i.e.

$$\rho(V_b^{\alpha}, V_b'^{\alpha}) = \rho(V_b^{\alpha}, V_d^{\alpha}) = p^{\alpha}.$$

Finally, since $V_b' = pV_d$, the correlation coefficient between $V_d^{\alpha}$ and $V_d'^{\alpha}$ is exactly the same as that between $V_b'^{\alpha}$ and $V_d'^{\alpha}$, i.e.

$$\rho(V_d^{\alpha}, V_d'^{\alpha}) = \rho(V_b'^{\alpha}, V_d'^{\alpha}) = \rho(V_b^{\alpha}, V_d^{\alpha}) = p^{\alpha}.$$

#### 4.2 Model with stochastic partitioning

We next focus on the correlation between birth and division sizes for the model with stochastic partitioning (model II). In this case, Eqs. (28) and (29) still hold and thus the key is to compute the variance of $V_b^{\alpha}$. Let $R = V_b'/V_d$ denote the partition ratio. Since $V_b' = RV_d$, we have

$$V_b'^{\alpha} = R^{\alpha}(V_b^{\alpha} + \Delta).$$

This shows that $R^\alpha(V_b^\alpha+\Delta)$ and $V_b^\alpha$ have the same distribution. Thus we obtain

$$\mathbb{E}V_b^\alpha = \mathbb{E}R^\alpha(V_b^\alpha+\Delta) = \mathbb{E}R^\alpha\mathbb{E}(V_b^\alpha+\Delta), \tag{35}$$

$$\mathbb{E}V_b^{2\alpha} = \mathbb{E}R^{2\alpha}(V_b^\alpha+\Delta)^2 = \mathbb{E}R^{2\alpha}\mathbb{E}(V_b^\alpha+\Delta)^2, \tag{36}$$

where we have used the fact that the partition ratio $R$ is independent of the birth size $V_b$ and generalized added size $\Delta$. Since $R$ has a beta distribution with mean $p$ and sample size parameter $\nu$, we have

$$\mathbb{E}R^\alpha = \frac{1}{B(p\nu,q\nu)}\int_0^\infty z^{\alpha+p\nu-1}(1-z)^{q\nu-1}dz = \frac{B(\alpha+p\nu,q\nu)}{B(p\nu,q\nu)}.$$

It then follows from Eq. (35) that

$$\mathbb{E}V_b^\alpha = \frac{(M_0+M_1)\mathbb{E}R^\alpha}{1-\mathbb{E}R^\alpha} = K_1(M_0+M_1), \tag{37}$$

where

$$K_1 = \frac{\mathbb{E}R^\alpha}{1-\mathbb{E}R^\alpha} = \frac{B(\alpha+p\nu,q\nu)}{B(p\nu,q\nu)-B(\alpha+p\nu,q\nu)}.$$

Similarly, it follows from Eq. (36) that

$$\mathbb{E}V_b^{2\alpha} = (2K_1+1)K_2(M_0+M_1)^2 + K_2\left[\frac{M_0^2}{N_0}+\frac{M_1^2}{N_1}\right], \tag{38}$$

where

$$K_2 = \frac{\mathbb{E}R^{2\alpha}}{1-\mathbb{E}R^{2\alpha}} = \frac{B(2\alpha+p\nu,q\nu)}{B(p\nu,q\nu)-B(2\alpha+p\nu,q\nu)}.$$

Combining Eqs. (37) and (38) shows that

$$\begin{aligned}\mathrm{Var}(V_b^\alpha) &= \mathbb{E}V_b^{2\alpha} - (\mathbb{E}V_b^\alpha)^2 \\ &= (2K_1+1)K_2(M_0+M_1)^2 + K_2\left[\frac{M_0^2}{N_0}+\frac{M_1^2}{N_1}\right] - K_1^2(M_0+M_1)^2 \\ &= \left[(2K_1+1)K_2-K_1^2\right](M_0+M_1)^2 + K_2\left[\frac{M_0^2}{N_0}+\frac{M_1^2}{N_1}\right].\end{aligned}$$

Inserting this equation into Eq. (28) finally shows that

$$\rho(V_b^\alpha,V_d^\alpha) = \sqrt{\frac{\left[(2K_1+1)K_2-K_1^2\right](M_0+M_1)^2 + K_2\left[\frac{M_0^2}{N_0}+\frac{M_1^2}{N_1}\right]}{\left[(2K_1+1)K_2-K_1^2\right](M_0+M_1)^2 + (K_2+1)\left[\frac{M_0^2}{N_0}+\frac{M_1^2}{N_1}\right]}}. \tag{39}$$

As $\nu\to\infty$, model II reduces to model I. In this case, we have

$$K_1 = \frac{p^\alpha}{1-p^\alpha},\quad K_2 = \frac{p^{2\alpha}}{1-p^{2\alpha}}.$$

Using these two equations, it is easy to check that $(2K_1+1)K_2-K_1^2=0$ and thus

$$\rho(V_b^\alpha,V_d^\alpha) = \sqrt{\frac{K_2}{K_2+1}} = \sqrt{p^{2\alpha}} = p^\alpha.$$

Therefore, the correlation coefficient given in Eq. (39) for model II reduces to the one given in Eq. (34) for model I.

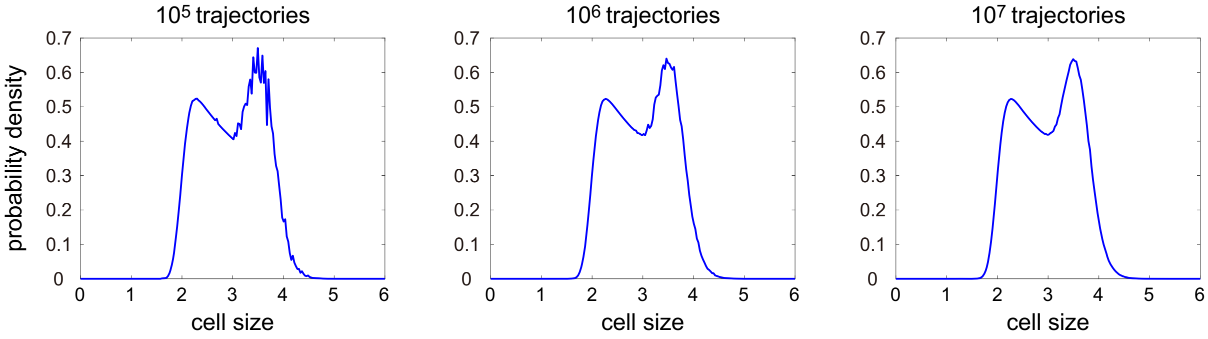


Fig. 1. **Cell size distributions obtained using stochastic simulations.** The three distributions are obtained by generating $10^5$, $10^6$, and $10^7$ stochastic trajectories, respectively. The parameters are chosen as $N = 50, r_0 = 0.6, r_1 = 0.1, g_0 = 0.01, g_1 = 2g_0, \alpha = 2, p = 0.5$. The parameters $a, M_0, M_1$ are chosen so that the mean cell size $\langle V \rangle = 3$.
